# Design of Allosteric Inhibitors for Mutant EGFR by Combined use of Machine Learning and Molecular Dynamics Simulations

**DOI:** 10.64898/2025.12.01.691623

**Authors:** Sapna Pal, Debasisa Mohanty

## Abstract

The non-small cell lung cancer (NSCLC)-associated Epidermal Growth Factor Receptor (EGFR) mutant L858R/T790M confers resistance to first- and second-generation tyrosine kinase inhibitors (TKIs). Allosteric inhibitors, binding outside the ATP-binding site, have emerged as alternative therapeutic agents. Unlike orthosteric inhibitors, they preferentially stabilize EGFR in an inactive conformation. Hence, understanding the mechanistic basis of this inhibition is essential for designing potent allosteric inhibitors. In this study, we performed microsecond-scale molecular dynamics (MD) simulations on the inactive conformations of apo-EGFR and apo-EGFR^L858R/T790M^ to explore how cancer-associated mutations induce a conformational switch toward the active kinase state. Simulations of allosteric inhibitor (EAI001) bound EGFR^L858R/T790M^ revealed that inhibitor binding enhances the inactive state population by disrupting the K745-E762 salt bridge and modulating key structural elements. These findings revealed the structural basis of allosteric inhibition in EGFR^L858R/T790M,^. It also emphasized the importance of MD simulations in allosteric drug design for assessing the ability of the inhibitor to enhance the population of the inactive state of the mutant EGFR. We have also standardized a virtual screening protocol involving screening of an allosteric TKI library by docking, re-ranking them with a machine learning-based scoring function (SG-ML-PLAP), and evaluating top-scoring molecules by MD and MM/GBSA to identify molecules stabilizing the inactive state of the EGFR^L858R/T790M^. This approach identified 10 novel allosteric kinase inhibitors predicted to be more potent than EAI001. Overall, our results not only elucidate the mechanism of allosteric inhibition in EGFR^L858R/T790M^ but also offer promising leads for the development of next-generation therapies to overcome TKI resistance.

## INTRODUCTION

EGFR is a transmembrane glycoprotein belonging to the ErbB family, which plays a critical role in regulating cell proliferation and differentiation by activating downstream signaling pathways through phosphorylation [1]. EGFR mutations are frequently observed in various cancers, including non-small cell lung cancer (NSCLC), head and neck cancer, breast cancer, and glioblastoma (GBM) [2–4]. These genomic aberrations are often associated with the continuous activation of signaling pathways, leading to uncontrolled cell proliferation and differentiation [2, 5]. Given the central role of EGFR kinase in cancer progression, multiple therapeutic strategies have been developed to target its activity, including small-molecule inhibitors, monoclonal antibodies, and antisense gene therapy [6–8]. Consequently, inhibiting EGFR kinase remains a crucial area of research, particularly in the development of novel targeted therapies.

At the structural level, the structure of EGFR consists of two lobes (N-lobe and C-lobe) and four primary domains: (1) extracellular ligand-binding domain, (2) transmembrane domain, (3) intracellular kinase domain, and (4) C-terminal regulatory domain. The activation of EGFR is initiated by the binding of a ligand, such as epidermal growth factor (EGF), to the extracellular domain. This ligand binding event triggers a series of allosteric change in the intracellular kinase domain, which subsequently induces EGFR dimerization [9]. The dimerized kinase domains can be homodimer which pairs with another EGFR, or heterodimer, which pairs with other proteins of ErbB family, such as ERBB2, ERBB3, and ERBB4. This dimerization event leads to trans-autophosphorylation of the kinase domain, catalyzing the transfer of a γ-phosphate from ATP to conserved tyrosine residues at the C-terminal of EGFR. This phosphorylation event triggers a cascade of downstream signaling events, leading to the activation of various signaling pathways, ultimately influencing cell survival, proliferation, and differentiation [10, 11].

Under physiological conditions, EGFR exists in a predominantly dimeric autoinhibited state and becomes activated upon the binding of a growth factor. These conformational transitions are governed by key structural motifs, including the phosphate-binding-loop (P-loop), activation-loop (A-loop), Asp-Phe-Gly (DFG)-motif, and αC-helix. The active conformation is characterized by the inward movement (“in”) of the αC-helix, an extended A-loop, and the formation of a critical salt bridge between K745 and E762, all of which stabilize ATP binding within the catalytic ATP-binding site. In contrast, the inactive conformation is marked by the outward movement (“out”) of the αC-helix, a collapsed A-loop, and a significant increase in the distance between the K745 and E762 residues to 14.4 Å [12, 13] (**Figure 1**).

**Figure 1.**
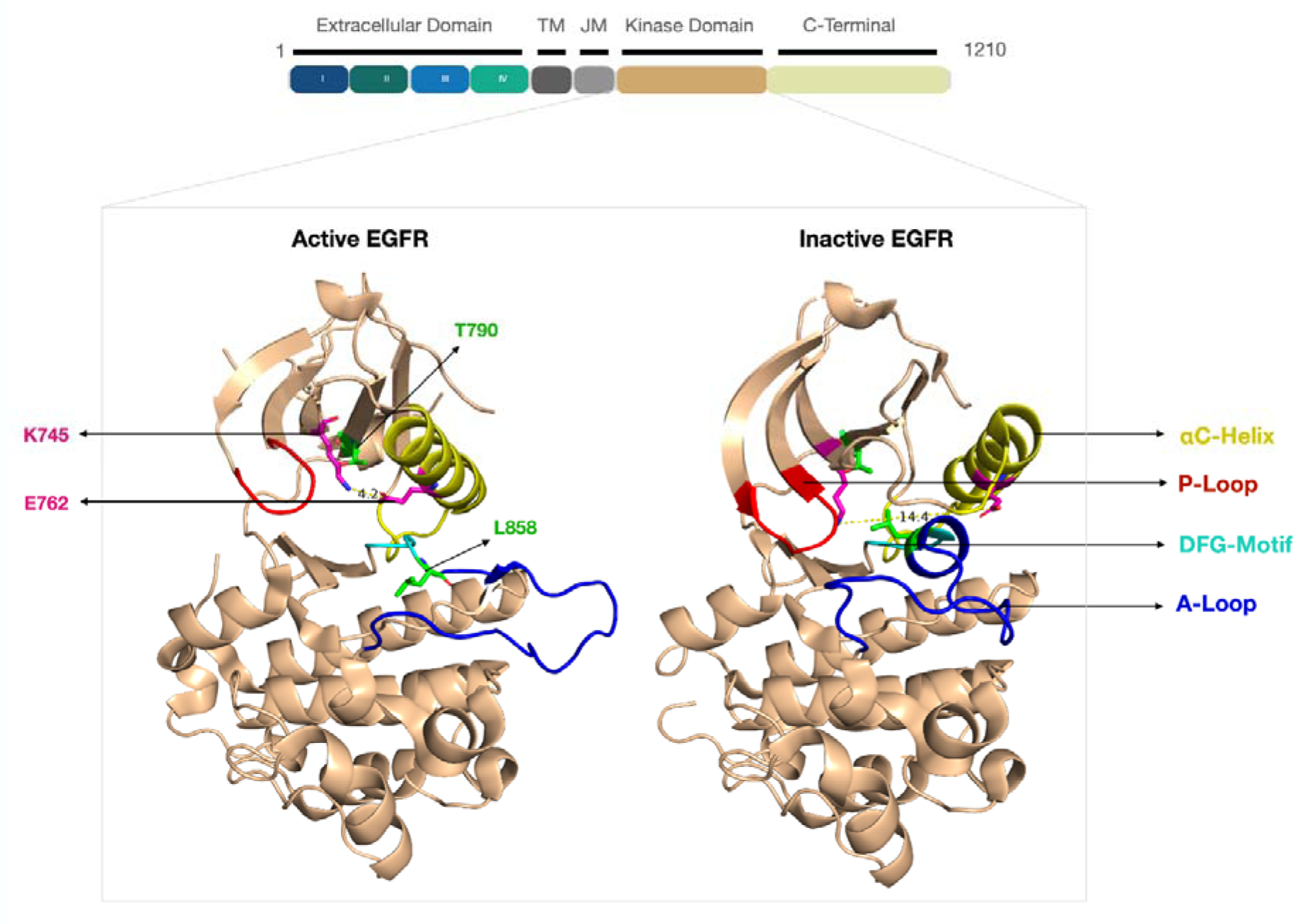
The EGFR kinase sequence with various domains, along with the crystal structure of active (PDB ID: 2GS2) and inactive (PDB ID: 4HJO) EGFR kinase domains, with various kinase substructures such as αC-helix, P-loop, DFG-motif, and A-loop, are shown in yellow, red, cyan, and blue colors, respectively. Diver/resistance mutation and K745-E762 salt bridge forming residue positions are shown in green and magenta color, respectively.

Crystallographic studies have significantly improved our understanding of structural transitions between active and inactive states of EGFR, as well as the potential intermediate conformations that exist between these states [14–17]. However, the static nature of crystallographic data has limitations in capturing the full extent of EGFR’s conformational dynamics. To address this, all-atom molecular dynamics (MD) simulations have been extensively employed to study the structural fluctuations and transitions of the EGFR kinase domain [12, 18, 19]. For instance, a study by Songtawee *et al.* analyzed the asymmetric movement of EGFR dimers in both active and inactive states, revealing a decrease in structural fluctuations in the active dimer, whereas no significant conformational changes were observed in the inactive dimer [19]. Another study, utilizing ∼50 microseconds of MD simulations, examined the active-to-inactive transitions of EGFR, identifying intermediate conformations that correlated with hydrogen-deuterium (H/D) exchange data, a technique used to investigate protein folding and structural dynamics [12]. MD simulations, in particular, allow for the detailed analysis of protein motions and interactions at the atomic level. However, the accuracy of MD simulations relies on careful selection of parameters, such as the non-bonded cutoff distance, which determines the range of interatomic interactions considered. An appropriate cutoff distance is crucial for capturing long-range electrostatic interactions, such as those involved in the K745-E762 salt bridge, which can significantly influence protein dynamics and function. Despite these advances, only a limited number of studies have leveraged microsecond- to millisecond-scale MD simulations to explore large-scale EGFR kinase domain folding and conformational transitions.

The abnormal structural transitions and stabilization of EGFR can also result from somatic mutations at critical residues. For example, L858, located within the A-loop, plays a crucial role in stabilizing the inactive EGFR conformation through hydrophobic interactions with N-lobe residues. Due to its functional importance, L858 is frequently mutated in NSCLC patients, leading to constitutive kinase activation [5]. Other commonly observed mutations include exon 19 deletions, kinase domain duplications, and rare mutations such as G719X, L861Q, and S768I [20, 21]. Various small-molecule kinase inhibitors have been designed to inhibit EGFR activation by targeting the ATP-binding site. However, prolonged use of these inhibitors frequently leads to the emergence of resistance mutations, the most notable being T790M. This secondary mutation in EGFR^L858R/T790M^ confers resistance to first-and second-generation ATP-competitive inhibitors that target the EGFR^L858R^. [22, 23].

To circumvent ATP-site resistance, non-ATP competitive allosteric inhibitors have been developed. These inhibitors bind to an allosteric site distinct from the ATP-binding pocket, inducing conformational changes that render the kinase domain inactive. One such inhibitor, EAI001, selectively targets the EGFR^L858R/T790M^ mutant, exhibiting an IC50 value of 24 nM at 1 mM ATP concentration[16]. The aminothiazole moiety of EAI001 interacts directly with the T790M mutation, which is hypothesized to contribute to its selectivity toward EGFR^L858R/T790M^ over wild-type EGFR. A derivative of EAI001, EAI045, was subsequently developed with improved binding affinity, although it lacks selectivity. Additionally, other allosteric inhibitors such as JBJ-04-125-02 and JBJ-07-149 have demonstrated promising efficacy, exhibiting high binding affinities to EGFR^L858R/T790M^ with IC50 values of 0.26 nM and 1.1 nM, respectively [24, 25]. Considering the importance of allosteric inhibitors, it is crucial to screen and develop new allosteric inhibitors with high binding affinity towards mutant EGFR^L858R/T790M^ which can change the conformation of EGFR as well. Given the therapeutic significance of allosteric inhibitors, ongoing research is essential to identify novel compounds with enhanced binding affinity for EGFR^L858R/T790M^. Additionally, the ability of these inhibitors to alter EGFR’s structural dynamics and favor its inactive conformation remains a critical area of investigation. Understanding these mechanistic insights will aid in the rational design of next-generation allosteric inhibitors that can effectively target drug-resistant EGFR mutants while minimizing off-target effects.

In this *in silico* study, we used all-atom (MD) simulations with an optimized non-bonded cutoff distance to accurately capture long-range interactions crucial for EGFR dynamics. We investigated the structural alterations in the inactive EGFR kinase domain caused by the L858R/T790M mutation and the impact of the allosteric inhibitor EAI001. Molecular dynamics simulations for 2µs on the inactive state structures of apo-EGFR^Wild^, apo-EGFR^L858R/T790M^, and EAI001-bound inactive EGFR^L858R/T790M^ revealed significant movements in critical substructures, including the αC-helix, A-loop, and P-loop. We analyzed the conformational landscape of EGFR^L858R/T790M^, its propensity for transition to the active state, and the effect of the allosteric inhibitor EAI001 on the dynamics of inactive to active state conformational transition. Finally, for the identification of potential allosteric inhibitors of EGFR^L858R/T790M^, we have standardized a virtual screening protocol involving the virtual screening of an allosteric kinase inhibitor library through docking, reranking their binding affinity with ML-based scoring function SG-ML-PLAP[26], and evaluation of the top candidates using MD simulations to analyze the structural impact of inhibitors on EGFR^L858R/T790M^ and the calculation of MM/GBSA binding free energy to assess their potential as novel therapeutic agents.

## MATERIALS and METHODS

### Preparation of EGFR kinase structures for simulation

The inactive state crystal structure (PDB ID: 5D41) of EGFR kinase with T790M mutation and bound to the allosteric inhibitor EAI001 was retrieved from the protein data bank [16]. The L858R mutation was modelled in the PDB structure 5D41 to prepare a double mutant inactive EGFR with a driver mutation L858R and a resistance mutation T790M. To generate wild type inactive EGFR model, the Met790 mutation was reverted to its original residue Thr790. All the mutations were created with the help of Chimera software[27]. As the activation loop region of the 5D41 crystal structure was not resolved, the loop region was modelled through the built-in Modeller extension in Chimera software. In the case of the active structure of EGFR kinase (PDB ID: 2GS2), two mutations L858R and T790M were modelled to create apo-active EGFR^L858R/T790M^ system. All the structures were modelled for the missing residues with the help of the Modeller tool[28] integrated in Chimera. Final systems for molecular dynamics (MD) simulations included: active apo-EGFR^Wild^, active apo-EGFR^L858R/T790M^, inactive apo-EGFR^Wild^, inactive apo-EGFR^L858R/T790M^ and inactive EAI001-EGFR^L858R/T790M^. The PDB structures, mutations and simulation time used in this study are provided in **Table 1**.

**Table 1.**
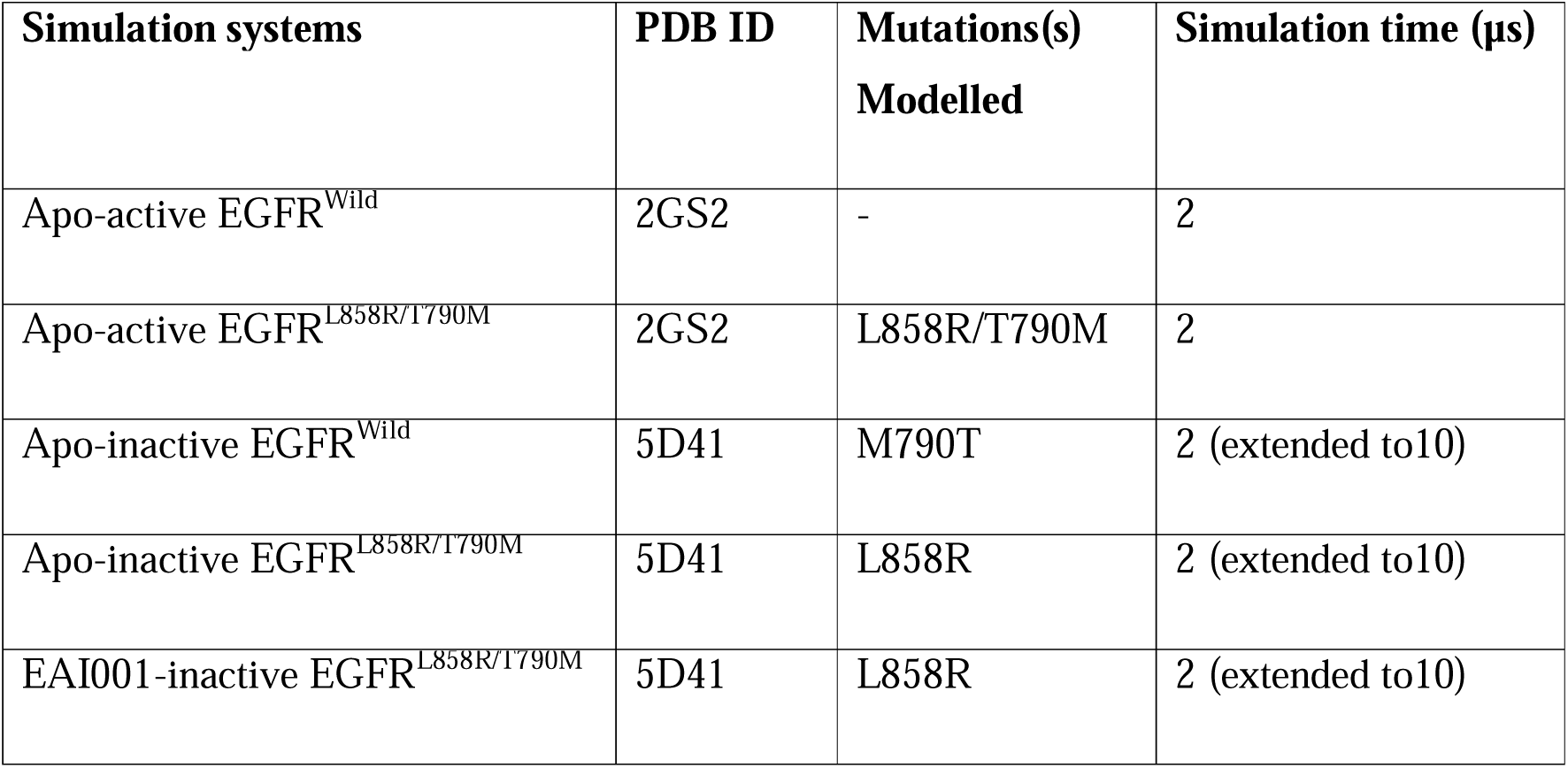
Information about the preparation of simulation systems, along with the simulation time.

### Preparation of ligand structures for simulation

The allosteric ligand EAI001 and other kinase allosteric inhibitors selected through virtual screening in this study were prepared by adding hydrogens and AM1-BCC charges[29]. Ligand parameters were generated using the General Amber Force Field (GAFF) via the Antechamber module [30, 31].

### Molecular Dynamics simulations

MD simulation for all three structures was carried out using the AMBER 20 [32] package and ff14SB force field[33]. In each simulation, the structures were solvated with explicit TIP3P water molecules using an octahedral water box that extended 10Å from the outermost protein atoms in all directions. The solvated system was neutralized by adding Na+ counter ions and additional Na+/Cl-were added to the box to achieve 150nM salt concentration. The solvated proteins were minimized for 5000 cycles with steepest descend and conjugate gradient. The restrain of 25 kcal mol^−1^ Å^−2^ was applied in the start of minimization and was reduced to 0 kcal mol^−1^ Å^−2^. After that the system was heated gradually to achieve the temperature of 300K over 100 ps of MD run in the NVT ensemble. During heating process the temperature was increased from 0 to 300K with the restrain of 10 kcal mol^−1^ Å^−2^. System was equilibrated in the NPT ensemble for 900ps of MD run with restrain gradually decreasing from 25 kcal mol^−1^ Å^−2^ to 0.1 kcal mol^−1^ Å^−2^. After this a final 5ns equilibration was done without restrain before starting the production step. A production dynamic was carried out without restrain for 2µs. Electrostatic interactions were calculated using the Particle Mesh Ewald (PME) method. Crucially, to ensure accurate capture of all relevant non-bonded interactions, especially considering the significant conformational changes between active (K745-E762 salt bridge distance typically < 4 Å) and inactive (K745-E762 salt bridge distance typically > 13 Å) states of EGFR, a non-bonded cutoff distance of 16 Å was employed.

### Analysis of molecular dynamics simulation trajectories

Analysis of MD trajectories was done in terms of 1) root mean squared deviation (RMSD) of backbone Cα-atoms, which measures the distance of atoms throughout the simulation with relation to the reference structure to analyze the overall stability and 2) root mean squared fluctuation (RMSF) of Cα-atoms, which measures the overall displacement of atoms throughout the simulation with relation to the reference structure to analyze flexible regions of protein structure. 3)The distance between salt bridge forming residues K745(NZ) and E762(CD) was also calculated throughout the 2µs production steps.

### Principal component analysis (PCA) analysis

Principal Component Analysis (PCA) was performed to analyze dominant protein motion patterns during the MD simulation production phase. PCA reduces the system’s dimensionality and identifies functionally relevant conformational changes. Analysis was conducted using cpptraj, and results were visualized using Matplotlib in Python[32, 34, 35].

### MM/GBSA for binding free energy calculations

The Molecular Mechanics Generalized Born Surface Area (MM/GBSA) method was used to estimate the relative binding free energy (ΔGbind) of EAI001 and selected allosteric inhibitors with EGFR^L858R/T790M^ (100ns) [36]. The binding free energy was calculated by extracting snapshots every 5 frames from the 100ns production step, resulting in a total of 20,000 frames (20ns) for the analysis using the following equation.

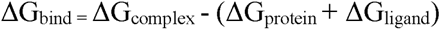

This equation is for calculating binding free energy (ΔG_bind_) difference between two states that is, bound state and free state of ligand and protein. Here, ΔG_bind_ represents relative binding free energy of the protein-ligand complex, ΔG_complex_ is free energy of the bound complex, ΔG_protein_ is free energy of unbound protein, and ΔG_ligand_ is free energy of unbound ligand.

### Virtual screening using docking simulations

#### Preparation of receptor

The minimized structure of the EAI001-bound inactive EGFR^L858R/T790M^ complex was used for docking studies. The EAI001 ligand was removed from the binding pocket to prepare the receptor for docking new ligands. The receptor was prepared for docking using AutoDock Tools, [37] which involves adding hydrogen atoms, assigning charges, and converting the structure into a format compatible with the docking software.

#### Preparation of chemical compound library

To screen allosteric inhibitors for inactive EGFR^L858R/T790M^, ChemDiv’s (https://www.chemdiv.com/) library of 26k allosteric kinase inhibitors was used for virtual screening. All the compounds were minimized using openbabel before preparing them for docking[38]. The ligands were prepared for docking using the prepare_ligand.py script from the ADFR suite [39], which converts the ligand structures into the appropriate format for AutoDock Vina.

#### Docking

For docking a grid was generated which was kept ligand centric, here the ligand was allosteric ligand i.e., EAI001 having coordinates surrounded by allosteric pocket. The re-docking of mutant EGFR^L858R/T790M^ - EAI001 complex was done for the validation of docking parameters. Finally, the grid size was kept 20×20x20 Å and the exhaustiveness was set to a default setting of 10. The docking was performed by AutoDock Vina and maximum of nine ligand poses were generated and the top-scoring pose was considered for further analysis [40].

#### SG-ML-PLAP re-scoring

The docked compounds on the allosteric site of EGFR^L858R/T790M^ were re-scored, and their binding affinity was predicted with the help of ML-based scoring function SG-ML-PLAP developed in our earlier work.

## RESULTS

### Dynamics of the wild-type EGFR starting from the active and inactive states

In order to understand the conformational dynamics of the EGFR, we carried out a 2µs MD simulation on the wild type EGFR starting from its active state crystal structure and modelled conformation for the inactive state. The model for the wild type inactive structure of EGFR was built using the crystal structure of inhibitor bound EGFR^T790M^ as a template. We calculated the root mean square deviation (RMSD) of the protein backbone to analyze the stability of the simulations. For both the active and inactive wild type EGFR, the RMSD exhibited higher fluctuations up to 1µs of the simulation time and stabilization was observed at the later stages of the simulations (**Supplementary Figure S1A**). The analysis of local flexibility of EGFR substructural regions using RMSF showed high fluctuations in the αC-helix region of the active EGFR structure, and on the other hand, the P-loop and A-loop regions displayed high fluctuations in the simulation of inactive EGFR, suggesting their role in the EGFR conformational state transition (**Supplementary Figure S1B**).

The critical structural feature that distinguishes the active from the inactive conformational state of EGFR is the formation of a salt bridge between the residues K745 and E762 within the kinase domain. Hence, we analyzed the distance between these salt bridge forming residues throughout the 2 µs simulation for both active and inactive EGFR structures. When starting from the active state, while the average salt bridge distance was 3.39Å, indicating its general formation, we observed its frequent fluctuations. Notably, within the initial 250 ns, the salt bridge distance transiently extended up to 7.5 Å, suggesting periods of instability. This dynamic behavior was observed throughout the entire 2µs simulation, with the salt bridge fluctuating from its stabilized states to exhibit frequent occurrences of conformational states where the K745-E762 distance extended to approximately ∼6.5 Å (**Figure 2**). In contrast, simulations initiated from the inactive wild type EGFR structure displayed a large distance between salt bridge forming residues, with an average of 14.1 Å (**Figure 2**), reflecting the absence of a salt bridge. Since the transition from inactive to active state and formation of salt bridge require large scale conformational changes, it was not possible to observe such large conformational transitions during the relatively short time scale of our simulation. Overall, time-series plots of the K745-E762 distance revealed the fluctuation distance from 3Å to 6.5Å in a few places in the active state, indicating formation and breakage of the salt bridge, while the salt bridge was completely absent in the simulations, which started from the inactive state. Complementary to these findings, and further illustrating the distinct conformational characteristics of the active/inactive state, analysis of the K745-E762 distance with a 13Å threshold (**Supplementary Figure S2, 1µs simulation**) showed that the active wild-type EGFR maintained the salt-bridge forming residue distance predominantly around 3-5Å, extending up to ∼6Å, whereas the inactive wild-type EGFR structure consistently displayed a distance of approximately 11Å.

**Figure 2.**
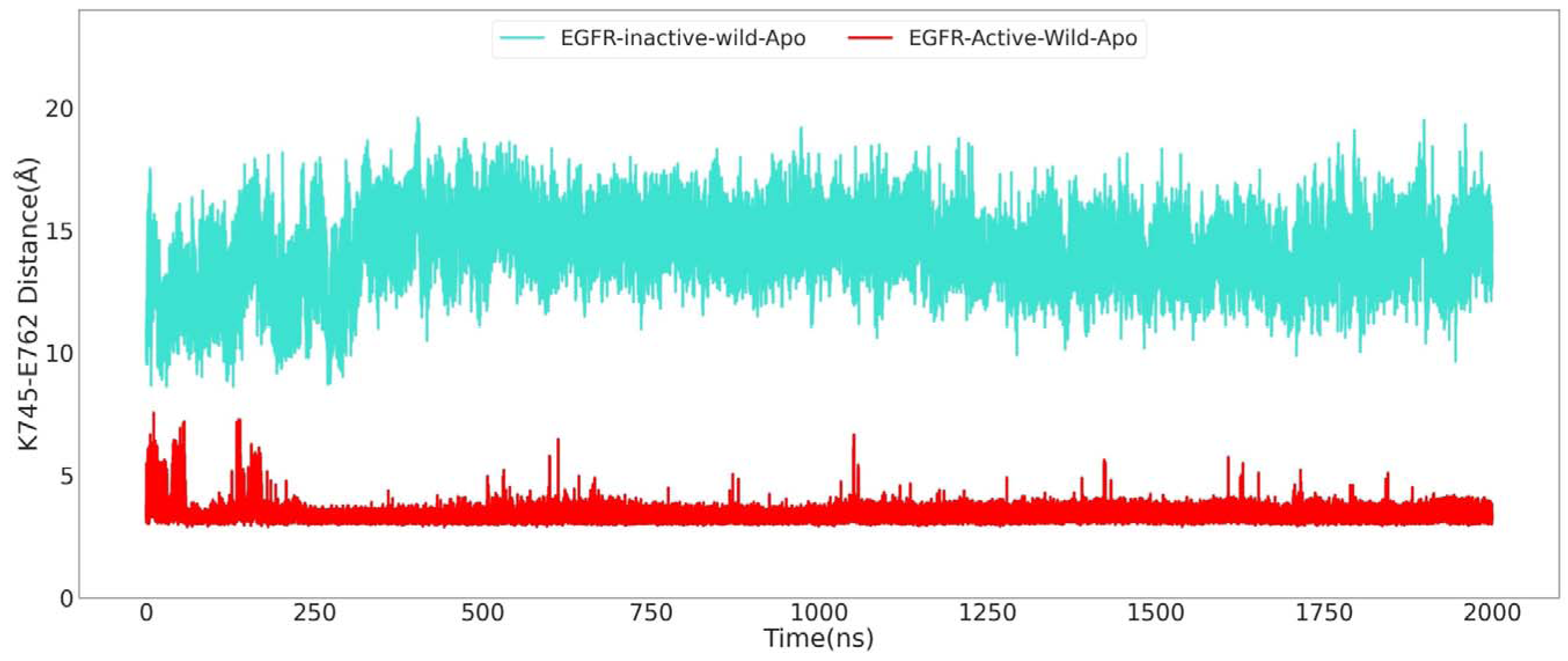
Time series plot of the distance between NZ and CD atom of K745 and E762 salt bridge forming residues, respectively. Distance calculated for the conformations observed during 2 µs simulations of active and inactive wild-type apo-EGFR.

### Dynamics of the double mutant EGFR^L858R/T790M^ starting from the active and inactive states

Following the characterization of wild type EGFR dynamics, we next investigated the L858R/T790M double mutant, which combines a common driver mutation (L858R) with a known resistance mutation (T790M). The overall stability of the mutant systems, analyzed through RMSD, showed relatively greater stabilization compared to their wild type counterparts (**Supplementary Figure S3A**). This suggests that the presence of these mutations contributes to a relatively stable conformational state. Analysis of the fluctuations in the local regions of EGFR further differentiated the mutant EGFR dynamics from wild type. Notably, in the case of mutant EGFR in the active state, we observed higher fluctuations across substructural regions, including αC-helix, A-loop, and at the latter part of the P-loop, as compared to inactive mutant EGFR (**Supplementary Figure S3B**).

Analysis of the distance between K745 and E762 revealed the presence of a highly stable salt bridge in the active state of the L858R/T790M double mutant, while the corresponding salt bridge was absent in the inactive state (**Figure 3**). For the active double mutant EGFR, an average distance of 3.31 Å was observed across the 2 µs simulation. Notably, unlike the frequent occurrences of conformations with K745-E762 distances extending up to 6.5Å seen in the active state, the mutant almost entirely eliminated such conformations, underscoring a significantly more constrained and persistently formed salt bridge. Conversely, for the inactive double mutant EGFR, the time-series plot of the distance between salt bridge forming residues across the 2 µs simulation showed its stabilization around ∼12 Å, with an average distance of 12.08 Å (**Figure 3**). While this still indicates an inactive-like state, analysis of the K745-E762 distance over the 2 µs trajectory revealed that the distance had reduced to ∼6Å at several time points, and the shift towards a slightly shorter average distance compared to the wild-type inactive (average 14.2 Å) indicated a conformational shift towards the active state. These results indicate that the double mutant is not only more stable in the active state compared to the wild type, unlike wild type EGFR, it also has a propensity for conformational switch towards the active state even when simulations were started from the inactive state. It is possible that in longer simulations inactive to active state conformational switch can be observed for the double mutant. These results provide a theoretical rationale for the pathogenic aberrant phosphorylation by the EGFR^L858R/T790M^ in NSCLC. Further analysis of the K745-E762 salt bridge distance with a 13Å threshold (**Supplementary Figure S4, 1µs simulation**) showed that the active L858R/T790M mutant structure showed sharp stabilization at an average distance of ∼3Å, while the inactive mutant structure displayed a highly dynamic and largely open conformation, qualitatively similar to wild-type inactive EGFR (**Supplementary Figure 1**).

**Figure 3.**
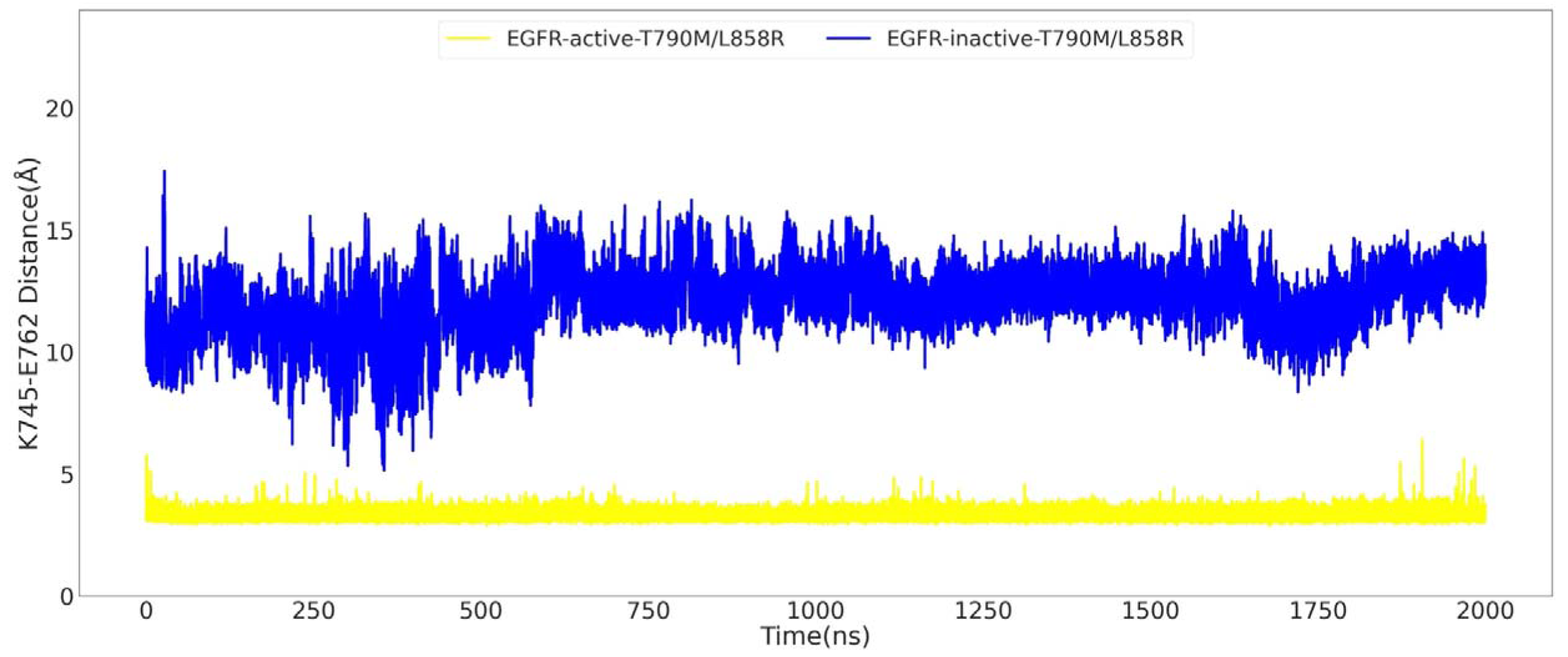
Time series plot of the distance between NZ and CD atom of K745 and E762 salt bridge forming residues, respectively. Distance calculated for the conformations observed during 2 µs simulations of active and inactive apo-EGFR^L858R/T790M^.

### Dynamics of EGFR^L858R/T790M^ bound to allosteric inhibitor EAI001

Since the double mutant has high propensity for aberrant phosphorylation and is resistant to orthosteric inhibitors which bind at the ATP binding site of the kinase, allosteric inhibitors like EAI001 have been developed. Allosteric inhibitors, which act by stabilizing an inactive enzyme conformation, represent a critical alternative therapeutic approach. Hence, we wanted to analyze the molecular basis of allosteric inhibition by carrying out 2µs MD simulation on the EAI001 bound inactive state structure of EGFR^L858R/T790M^ double mutant. The K745-E762 salt bridge is crucial for stabilizing the active conformation of the EGFR kinase domain and facilitating ATP binding. Hence, the distance between these two salt bridge forming residues was used as reaction coordinate to analyze inactive to active state conformational transition of the EAI001 bound double mutant. **Figure 4A** shows the overlapped density plots for the distribution of K745-E762 distances observed for the inactive apo-EGFR^Wild^, inactive apo-EGFR^L858R/T790M^, and inactive EAI001-EGFR^L858R/T790M^. As explained earlier, in wild type, the average distance was 14.2Å and minimum of 10Å indicating a stable inactive conformation. In contrast, the apo−EGFR^L858R/T790M^ mutant showed a decreased average distance of ∼12 Å and minimum of 7.5Å, suggesting that the mutations promote a shift towards an active-like conformation. However, the EAI001-bound EGFR^L858R/T790M^ exhibited a more complex behavior, with three distinct peaks in the salt bridge distance distribution: a small peak at ∼9.5 Å, a large peak at ∼12 Å, and another peak at ∼14 Å. This suggests that the inhibitor modulates the dynamics of the mutant EGFR kinase, inducing a conformational shift towards the inactive state (**Figure 4A**). Furthermore, extending the simulation to 10 µs and analyzing the last 2 µs timeframe (8-10 µs) (**Figure 4B**) revealed a clearer dominant peak around ∼14 Å for the EAI001-bound EGFR^L858R/T790M^, indicating that over longer timescales, the inhibitor stabilizes inactive conformation. These findings highlight the complex interplay between the L858R/T790M mutation, the allosteric inhibitor EAI001, and the dynamics of the K745-E762 distance. The mutation appears to promote a shift towards the active conformation, while the inhibitor counteracts this effect by destabilizing the salt bridge and hindering its formation as evidenced by the longer timescale stabilization around ∼14 Å. This detailed analysis provides valuable insights into the molecular mechanisms underlying the activation of EGFR and the inhibitory action of EAI001.

**Figure 4.**
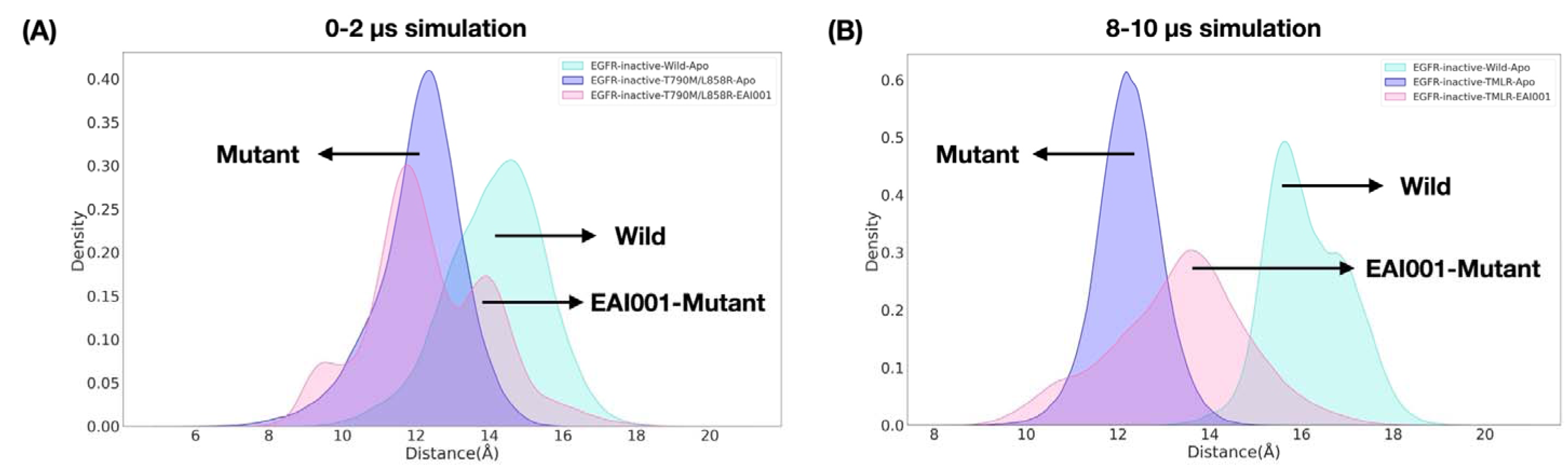
Salt bridge distance between NZ and CD atom of K745 and E762, respectively. Distribution of salt bridge distance in conformations observed during 0-2 and 8-10 µs simulations of wild-type apo-inactive EGFR, apo-inactive EGFRT790M/L858R, and EAI001-inactive EGFRT790M/L858R (A, B).

Next, to analyze various substructural dynamics of the kinase domain we employed Principal Component Analysis (PCA) to identify the major collective motions of the protein during the simulations. **Figure 5** shows the PCA plot, where the yellow regions in the plot indicate clusters of similar conformations sampled during the simulations. We extracted representative structures from the highly populated clusters in each simulation system (wild-type, apo-mutant, and inhibitor-bound mutant) to compare the conformations of key substructures (P-loop, A-loop, DFG-motif, and αC-helix) across these representative structures, and gain insights into how the L858R/T790M mutation and the allosteric inhibitor influence the conformational dynamics of the EGFR kinase domain.

**Figure 5.**
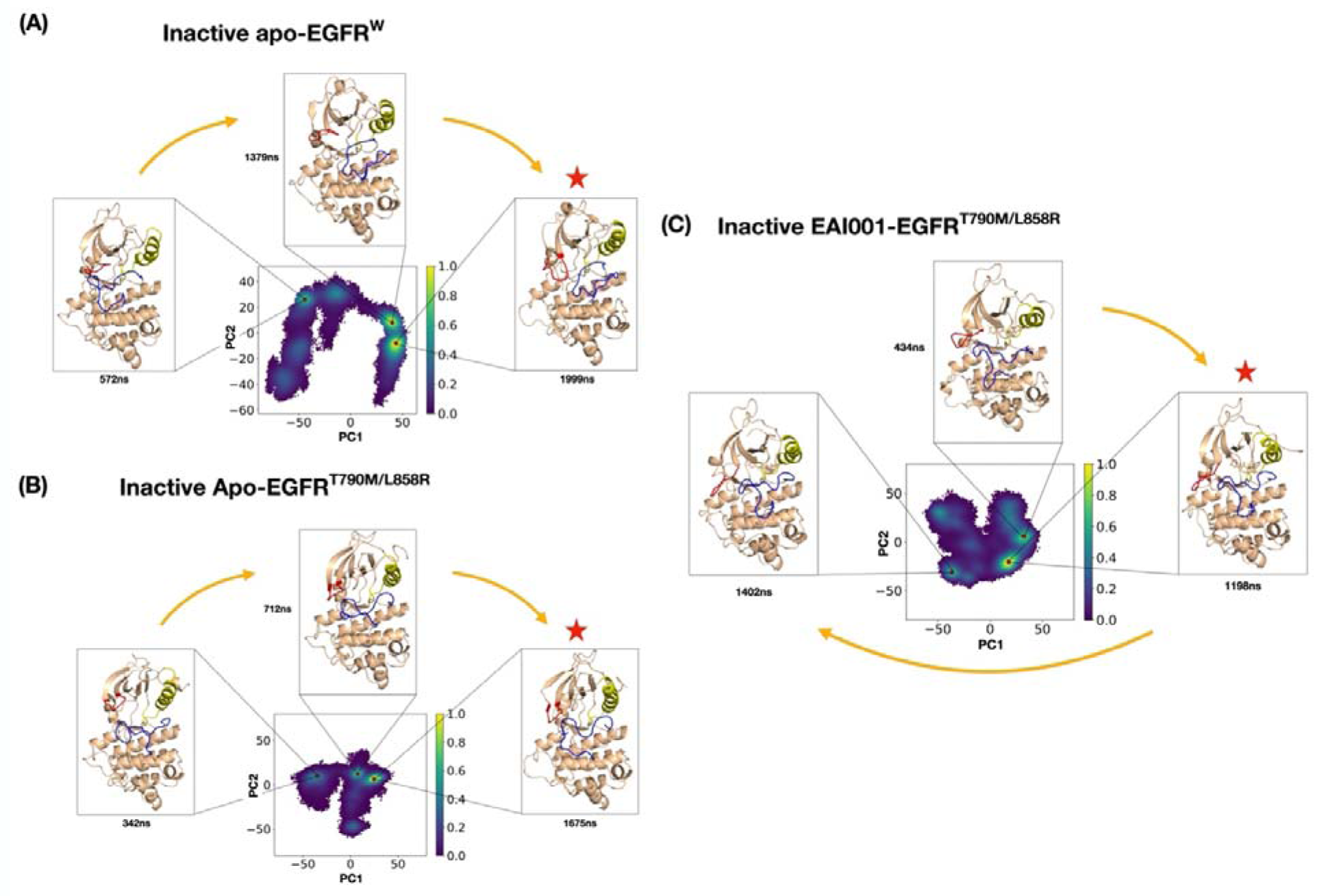
Energy landscape projection created by Principal Component Analysis (PCA) to analyze the greatest variance in the movement of structures over the simulation. The plot utilizes the first two eigenvectors. The red dot represents the conformation extracted from a highly populated cluster in inactive apo-EGFR^Wild^ (A), apo-EGFR^T790M/L858R^ (B), and EAI001-EGFR^T790M/L858R^ (C).

Analysis of the αC-helix, a key structural element involved in kinase activation, revealed distinct movements across different systems. In the inactive apo-EGFR^Wild^, the three major conformations obtained from the PCA clusters at 572ns, 1379ns, and 1999ns showed no significant movement of the αC-helix (**Figure 5A**). This observation was further supported by the overlap of the conformation from the highest PCA cluster with the reference crystal structure 5D41 with modelled A-loop (**Figure 6A**). Interestingly, in the inactive apo-EGFR^L858R/T790M^, a significant inward movement of the αC-helix was observed as the simulation progressed, suggesting a shift towards an active-like conformation (**Figure 5B**). This inward movement was also evident in the structural overlap with the reference crystal structure (**Figure 6A**). In contrast, for the EAI001-bound inactive EGFR^L858R/T790M^, a significant loss of the starting αC-helix was observed in the majority of the population, as seen in the structure at 1198ns (**Figure 5C and 6A**). However, a small cluster formed at the later stage of the simulation exhibited reformation of the αC-helix (**Figure 5C)**.

**Figure 6.**
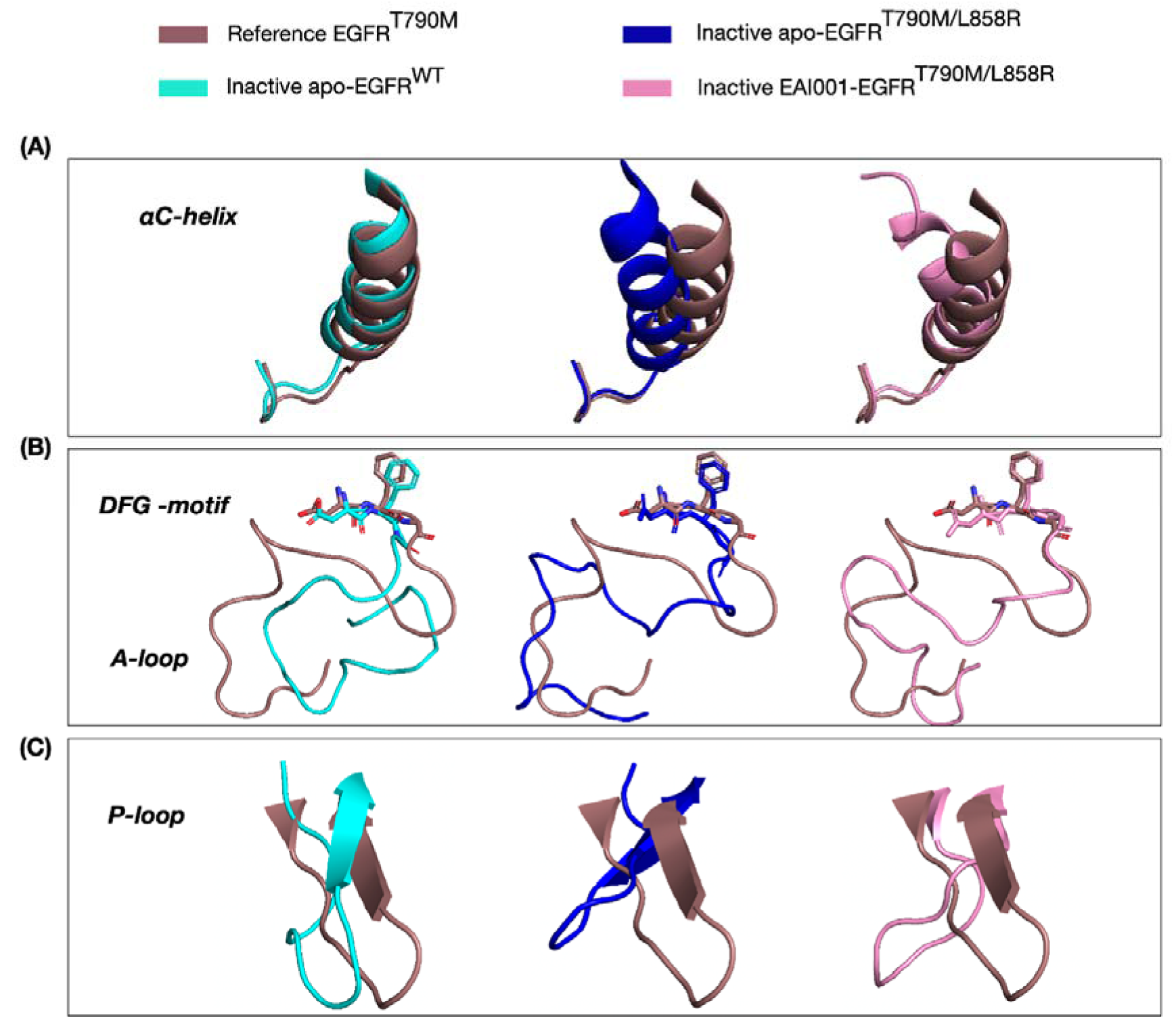
Superimposition of the EGFR kinase conformations showing αC-helix (A), DFG-motif and A-loop (B), and P-loop (C) (taken from the highly populated PCA cluster) onto the inactive EGFR crystal structure (PDB ID: 5D41) taken as a reference.

To further analyze the changes in the secondary structure of the αC-helix, we calculated the alpha-helix occupancy of the inactive EAI001-EGFR^L858R/T790M^ throughout the simulation. The results (**Supplementary Figure S5**) clearly showed the decrease in the alpha-helix occupancy at the start of the αC-helix upon inhibitor binding, suggesting the formation of a local disordered state in the mutant EGFR [12, 41]. This observation is consistent with previously reported inactive EGFR structures with shorter αC-helices (PDB ID: 2GS7, 4HJO).

The A-loop, another crucial regulatory element, was modelled in our simulations due to missing coordinates in the inactive crystal structure. The MD simulations provided valuable insights into its flexibility. Our analysis revealed high flexibility of the modelled A-loop in all three systems. In the inactive apo-EGFR^Wild^, the A-loop contracted in the center (**Figure 5A and 6B**), potentially representing an intermediate position before forming a short helix at its N-terminal and blocking the ATP-site. A similar, albeit less pronounced, contraction was observed in the EAI001-bound EGFR^L858R/T790M^ (**Figure 5C and 6B**). In contrast, the A-loop of the apo-EGFR^L858R/T790M^ mutant was more relaxed and extended towards the left-hand side (**Figure 5B and 6B**). A short α-helix formed at the N-terminal portion of the A-loop (Gly857–Gly863) is a key element that prevents the EGFR kinase from being active until the ligand binds and induces the conformational change [41]. Since no α-helix formation was observed at the A-loop in any of the systems, we analyzed the changes in its secondary structure. While no helix formation was observed, an increase in turn like conformation (prior to formation of helix) was observed in both the inactive apo-EGFR^Wild^ and the EAI001-EGFR^L858R/T790M^ compared to the apo-EGFR^L858R/T790M^, suggesting an increased propensity for helix formation (**Supplementary Figure S6**). This was confirmed by the formation of an α-helix at the end of an 8.5µs simulation of the EAI001-bound EGFR^L858R/T790M^ (**Supplementary Figure S7**), which was not observed in the other two systems even after extending the simulations to 10µs. No significant movement of the DFG-motif was observed in any of the simulation systems (**Figure 6B**).

In the inactive EGFR state, the P-loop is folded inwards and interacts with the αC-helix through hydrophobic interactions, maintaining the αC-helix-out conformation [42]. Our simulations revealed that the P-loop in the wild-type inactive EGFR was slightly coiled and moved inwards relative to the apo-EGFR^L858R/T790M^ and EAI001-EGFR^L858R/T790M^. The P-loop in the EAI001-bound system was not folded inwards to the same extent, with its movement falling somewhere between the apo-EGFR^L858R/T790M^ and apo-EGFR^Wild^ (**Figure 6C**). This suggests that both the mutation and the inhibitor can influence the conformation and dynamics of the P-loop, potentially affecting its interaction with the αC-helix and the overall stability of the inactive state.

Lastly, we also analyzed the distance between salt bridge forming residues in representative conformations extracted from the most populated PCA clusters (**Supplementary Figure S8**). The largest distance (14.4Å) was observed in the wild-type, followed by 13.9Å in the EAI001-bound mutant. The apo-mutant showed the shortest distance (11.2Å), reinforcing the notion that the mutation favors a conformation closer to the active state. The intermediate distance in the EAI001-bound mutant, along with the multi-modal distribution, suggests that the inhibitor induces a more dynamic behavior of the salt bridge, preventing it from adopting the active conformation.

The development of conventional TKIs depends upon how strongly they bind to the kinase orthosteric pocket to carry out competitive inhibition against ATP. However, the EGFR allosteric inhibitors work by altering the conformation of mutant EGFR^L858R/T790M^ and stabilizing the inactive conformational state. Hence, for the development of a better allosteric inhibitor, it is crucial to analyze its ability to prevent the shift of EGFR^L858R/T790M^ into an active like state along with the high binding affinity to the allosteric pocket. Therefore, we developed a virtual screening protocol where high affinity kinase allosteric binders are screened through molecular docking studies, followed by MD simulations to analyze their effect on the dynamics of EGFR^L858R/T790M^ to stabilize it in an inactive conformational state.

### Virtual screening of allosteric kinase inhibitors with mutant EGFR^L858R/T790M^ kinase revealed top allosteric binders

In order to identify novel allosteric inhibitors with potentially higher affinity for the EGFR^L858R/T790M^ mutant than the known inhibitor EAI001, we conducted a virtual screening. For that, we took the minimized conformation of inactive-EGFR^L858R/T790M^, which was minimized with EAI001 to adjust the inhibitor in the pocket of L858R/T790M mutant EGFR. Before screening a library of compounds, we first validated our docking protocol by re-docking EAI001 into the allosteric pocket of EGFR^L858R/T790M^ using AutoDock Vina. The re-docked ligand pose closely resembled the original binding pose observed in the minimised structure (**Supplementary Figure S9A**), confirming the reliability of our docking parameters. The predicted binding free energy for EAI001 was −9.9 kcal/mol. Analysis of the re-docked complex revealed key interactions between EAI001 and the allosteric binding pocket residues (**Supplementary Figure S9B**). These interactions include hydrogen bonds with Asp855 and Lys745, and hydrophobic interactions with Met790, Leu788, Met766, Leu777, Ile759, Glu762, and Ala763. Importantly, this interaction pattern is consistent with that observed in the minimized EAI001-EGFR^L858R/T790M^ complex and crystal structure of the EAI001-EGFR^T790M^ complex, further validating our docking approach. With a validated docking protocol, we proceeded to screen a library of allosteric kinase inhibitors to identify promising candidates that could bind to the allosteric site of the EGFR^L858R/T790M^ mutant with high affinity.

A library of 26,318 kinase allosteric inhibitors from ChemDiv were docked in the allosteric site of the EGFR^L858R/T790M^ kinase domain (**Figure 7A**). Since the docking scores often do not show good correlation with the experimental binding affinities, we used an ML-based scoring function SG-ML-PLAP to refine our selection and improve the accuracy of binding affinity predictions. In our earlier work, SG-ML-PLAP was trained and validated on a large dataset of protein-ligand complexes with known binding affinities [26]. It was found that, out of the 26,318 compounds, 12961 had SG-ML-PLAP scores above that of EAI001. From this list of top scoring compounds, we selected the top ten compounds exhibiting higher predicted binding affinities than EAI001. These compounds showed binding energy scores ranging from 9.61 to 9.27, compared to 6.32 for EAI001, suggesting stronger interactions with the allosteric pocket. The Autodock VINA and SG-ML-PLAP scores of these selected compounds are listed in **Table 2**. As intended, all ten compounds were predicted to bind within the allosteric pocket of EGFR (**Figure 7B**). To further assess the suitability of these compounds as potent allosteric inhibitors of EGFR^L858R/T790M^, we evaluated their drug-likeness and physiochemical properties. The radar plot (**Supplementary Figure S10**) generated by ADMETLAB3.0 for the predicted properties revealed that all ten compounds fell within the acceptable range, indicating favorable drug-like characteristics[43]. This suggests that these compounds not only exhibit high affinity for the EGFR^L858R/T790M^ allosteric site but also possess promising drug-like properties, warranting further investigation as potential lead compounds for drug development.

**Figure 7.**
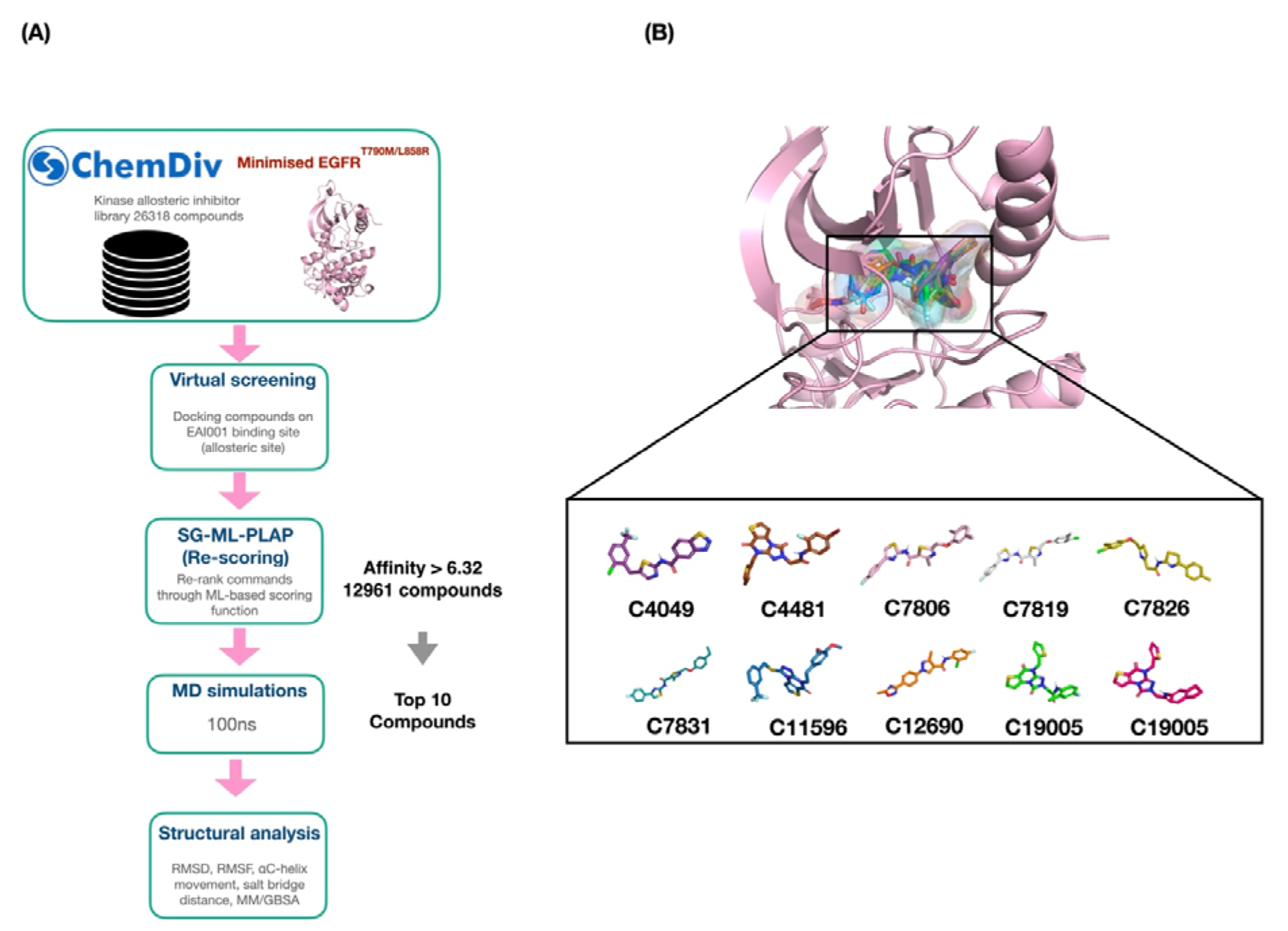
Virtual screening pipeline for finding the potential allosteric inhibitors for EGFR^L858R/T790M^ (A). The top ten compounds selected for simulations with the binding affinity (SG-ML-PLAP score) higher than EAI001 (B).

**Table 2.** AutoDock Vina scores and SG-ML-PLAP predicted score for the binding affinity of EAI001 and top ten compounds obtained after docking.

| **Name** | **SG-ML-PALP score** | **Docking Score (kcal/mol)** |
| --- | --- | --- |
| C4481 | 9.61 | -7.2 |
| C19005 | 9.54 | -6.4 |
| C11596 | 9.34 | -7.4 |
| C7806 | 9.34 | -6.5 |
| C7826 | 9.34 | -7.0 |
| C7831 | 9.34 | -6.1 |
| C7819 | 9.32 | -5.9 |
| C12690 | 9.31 | -4.7 |
| C4049 | 9.30 | -7.3 |
| C19017 | 9.27 | -6.7 |
| EAI001 | 6.32 | -9.9 |

To further evaluate the top ten allosteric inhibitors identified through virtual screening, we performed 100ns MD simulations for each compound in complex with the inactive EGFR^L858R/T790M^ kinase. This allowed us to assess the stability of the complexes and analyze the impact of the inhibitors on the EGFR^L858R/T790M^ kinase dynamics. Analysis of the RMSD of both the protein and the ligands revealed that most complexes were stable throughout the 100ns simulations (**Supplementary Figure S11A and B**). All compounds, except for C7806 and C7826, showed average RMSD values within 2.5Å, indicating stable binding. The EGFR^L858R/T790M^ kinase itself also remained stable in all simulations, with average RMSD values within 3Å, except for C7806. Next, we analyzed the RMSF values of the EGFR^L858R/T790M^ residues simulated with allosteric inhibitors to understand the atomic fluctuation of substructural regions. C7806, C7831, C7826, and C4481 showed high fluctuation in αC-helix, C11596, C19005, C7826, and C12690 showed high fluctuation in A-loop, and C12960, C19017, and C19005 showed high fluctuation on P-loop (**Supplementary Figure S11C**). The superimposition of the final conformations of EGFR^L858R/T790M^ after 100ns simulations with the initial minimized structure (**Supplementary Figure S12**) revealed significant structural changes in αC-helix induced by some of the compounds. Notably, C7806, C4049, and C19005 caused a pronounced outward movement of the αC-helix, even greater than that observed with EAI001. Also, C7831 led to a reduction in the αC-helix. Overall, structural changes caused by allosteric inhibitors in the mutant EGFR^L858R/T790M^ were observed in the majority of cases.

We also analyzed the impact of the inhibitors on the K745-E762 distance, a key determinant of EGFR activation. Notably, all compounds exhibited a significantly larger average distance (ranging 11.8 Å to 15.9 Å) between salt bridge forming residues when bound to the inactive EGFR^L858R/T790M^ compared to EAI001-bound inactive EGFR^L858R/T790M^ during the 100ns simulation. The compound C7806, C4049, C7831, and C12690 showed the most significant increases, followed by C4481, C19017, C19004, C11596, C7826, and C7819 (**Figure 8**). These short 100 ns simulations suggest that the screened compounds hold promise as potential allosteric inhibitors of EGFR^L858R/T790M^ by stabilizing the inactive state. However, longer timescale simulations will be required to fully characterize the long-term effects of these compounds on the EGFR conformation and their inhibitory mechanisms.

**Figure 8.**
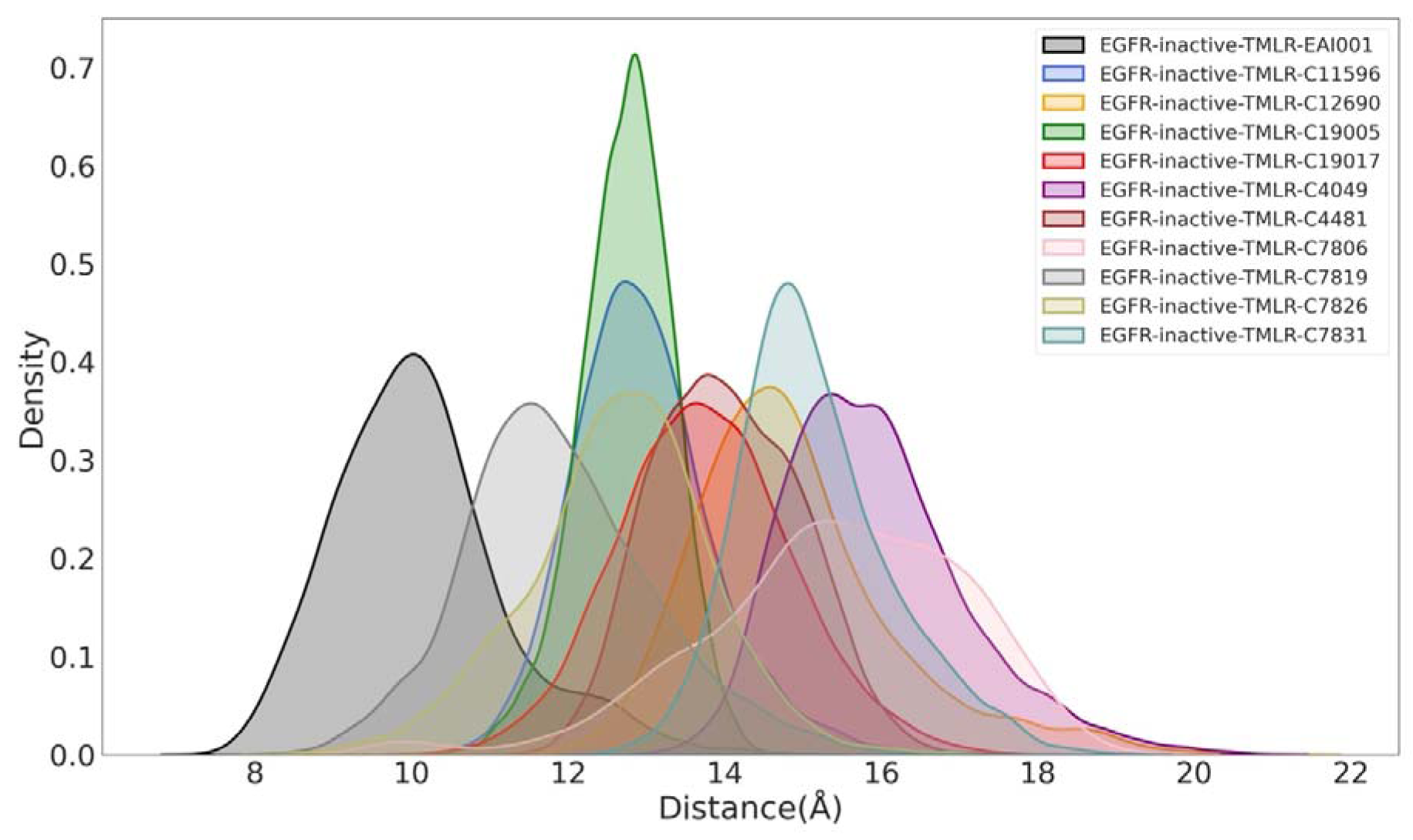
Density plot of EGFR^L858R/T790M^ conformations with distance between the salt-bridge forming residues NZ and CD atom of K745 and E762, respectively. The apo-inactive EGFR^L858R/T790M^ conformations were simulated with selected top ten compounds and EAI001 for 100 ns.

Next, we calculated the relative binding free energies (ΔG^total^) of the compounds using MM/GBSA analysis (**Table 3**). Interestingly, all compounds, including the top three C11596 (−50.76 kcal/mol), C7831(−49.52 kcal/mol), and C4481(−49.83 kcal/mol), showed higher binding affinities than EAI001 (−35.12 kcal/mol). Finally, a multi-criteria approach was employed to select the best binder candidates. **Table 4** outlines the categories used, including docking score, ML-based predicted score, αC-helix displacement, salt-bridge distance, and MM/GBSA scores. Compounds were selected if they ranked within the top five in at least three of these categories, ensuring a comprehensive assessment of their potential. Taken together, these results indicate that compounds C4481, C7806, C4049, C11596, and C7831 are promising candidates for further development as allosteric inhibitors of EGFR^L858R/T790M^. They exhibit high binding affinities, induce significant structural changes that favor the inactive conformation, and possess favorable drug-like properties.

**Table 3.** MM/PBSA analysis for calculating the binding affinity of EAI001 and top ten compounds with EGFRL858R/T790M after simulations.

|  | **VDW** | **EEL** | **EGB** | **ESURF** | **ΔG GAS** | **ΔG SOLV** | **ΔG TOTAL** |
| --- | --- | --- | --- | --- | --- | --- | --- |
| **C11596** | -68.35 | -19.64 | 45.48 | -8.25 | -87.99 | 37.23 | -50.76 |
| **C7831** | -66.32 | -35.91 | 60.01 | -7.70 | -102.23 | 52.31 | -49.92 |
| **C4481** | -69.80 | -8.29 | 35.49 | -7.27 | -78.10 | 28.21 | -49.83 |
| **C4049** | -60.42 | -50.05 | 69.13 | -7.13 | -110.48 | 62.0 | -48.47 |
| **C7806** | -66.0 | -29.38 | 56.59 | -8.10 | -95.38 | 48.49 | -46.89 |
| **C19005** | -63.78 | -14.65 | 38.84 | -7.21 | -78.43 | 31.62 | -46.81 |
| **C7819** | -59.16 | -15.-5 | 36.62 | -7.52 | -74.22 | 29.10 | -45.12 |
| **C19017** | -59.20 | -12.19 | 37.96 | -6.82 | -71.36 | 31.14 | -40.21 |
| **C12690** | -52.66 | -4.27 | 25.85 | -6.58 | -56.93 | 19.26 | -37.66 |
| **C7826** | -53.27 | -9.75 | 32.20 | -6.82 | -63.03 | 25.38 | -37.65 |
| **EAI001** | -49.54 | -17.52 | 37.99 | -6.06 | -67.06 | 31.93 | -35.12 |

**Table 4.** MM/PBSA analysis for calculating the binding affinity of EAI001 and the top ten compounds with EGFRL858R/T790M after simulations.

| **Name** | **SG-ML-**<br>**PLAP**<br>**score rank** | **αC-helix**<br>**displacement**<br>**rank** | **salt-bridge**<br>**distance**<br>**rank** | **MM/GBSA**<br>**scores rank** | **Docking**<br>**score rank** |
| --- | --- | --- | --- | --- | --- |
| **C4481** | **1** | **6** | **5** | **3** | **4** |
| C19005 | 2 | 3 | 8 | 6 | 8 |
| **C11596** | **3** | **8** | **7** | **1** | **2** |
| **C7806** | **4** | **1** | **1** | **5** | **7** |
| C7826 | 5 | 11 | 9 | 10 | 5 |
| **C7831** | **6** | **4** | **3** | **2** | **9** |
| C7819 | 7 | 10 | 10 | 7 | 10 |
| C12690 | 8 | 9 | 4 | 9 | 11 |
| **C4049** | **9** | **2** | **2** | **4** | **3** |
| C19017 | 10 | 7 | 6 | 8 | 6 |
| EAI001 | 11 | 5 | 11 | 11 | 1 |

## DISCUSSION

EGFR kinase inhibitors play a crucial role in targeting mutant EGFR-driven cancers; however, the emergence of drug resistance remains a significant challenge. Extensive research efforts have been dedicated to developing novel inhibitors against drug-resistant EGFR mutations. In this study, we employed all-atom MD simulations to investigate the conformational changes associated with the L858R/T790M double mutation in inactive EGFR. By utilizing a high-range non-bonded interaction cutoff of 16 Å, we were able to capture key structural shifts that differentiate mutant EGFR from its wild-type counterpart. Our simulations revealed that wild-type EGFR in its inactive state adopts a conformation, characterized by a stabilized outward-positioned αC-helix, and a potentially contracted A-loop. Upon introducing the L858R/T790M double mutation, a distinct shift toward an active-like conformation was observed, marked by the inward movement of the αC-helix and an extended A-loop. Although complete formation of the stable K745-E762 salt bridge was not observed for EGFR^L858R/T790M^, when simulations were started from the inactive conformation, our extended simulation analysis indicated a decrease in the distance between salt-bridge-forming residues upon mutation. This structural shift suggests that the L858R/T790M mutation enhances the propensity of kinase toward active state, potentially contributing to drug resistance mechanisms. Furthermore, our study provides mechanistic insights into the allosteric inhibition of mutant EGFR by EAI001. Simulations of L858R/T790M mutant EGFR in complex with EAI001 revealed the adoption of an intermediate conformation, featuring a disordered αC-helix, a partially contracted A-loop, and a larger distance between salt bridge forming residues. These conformational changes indicate that in our simulations, EAI001 could not induce complete transition to the inactive state of EGFR^L858R/T790M^, with the potential for increased stabilization over extended simulation time. Overall, our simulations bridge this knowledge gap by elucidating how the driver/resistance mutation alters structural response of EGFR to allosteric inhibition.

In search of novel and more effective allosteric inhibitors, we conducted virtual screening using the ChemDiv kinase allosteric inhibitor library. The docked complexes were reranked using SG-ML-PLAP, a recently developed ML based scoring function. The top 10 compounds were selected based on their SG-ML-PLAP scores, and subsequently, 100 ns MD simulations were performed to assess their potential to induce the transition of EGFR^L858R/T790M^ to the inactive state by binding to the allosteric site. Binding free energy calculations using MM/GBSA analysis further validated the high affinity of these compounds for the mutant kinase. Among the screened molecules, by using muti-criteria approach, compounds C4481, C7806, C4049, C11596, and C7831 demonstrated superior allosteric binding compared to EAI001, making them promising candidates for further investigation. Considering the scarcity of highly selective allosteric inhibitors for EGFR^L858R/T790M^, our study provides crucial insights into potential high-affinity compounds that could serve as next-generation allosteric inhibitors. Future *in vitro* and *in vivo* studies will be necessary to validate their efficacy and further explore their potential therapeutic applications. In summary, in the current work, we have developed a novel *in silico* protocol for the identification of allosteric inhibitors that can alter the dynamics of conformational transitions.

## Data Availability

The data that supports the findings of this study are available at https://www.rcsb.org/ and in the supplementary material of this article.

## Supporting Information

Supplementary_Figures.pdf: Supplementary Figures S1 to S12

## Author Contributions

**Sapna Pal**: Methodology, Investigation, Data curation, Validation, Writing - original draft.

**Debasisa Mohanty**: Conceptualization, Supervision, writing (review and editing), funding acquisition.

## Acknowledgments

This work was supported by the Department of Biotechnology, Government of India grant to National Institute of Immunology, New Delhi. DM acknowledges financial support from Department of Biotechnology (DBT), India under R&D project grants (BICB: BT/PR40325/BTIS/137/1/2020), (DIAU: BT/BI/TCB/007/2021) and (NNP: BT/PR40267/BTIS/137/67/2023 & BT/PR40160/BTIS/137/64/2023). The work was also supported by National Supercomputing Mission, MeiTY, India under NSM-NPGDD project (MeitY/R&D/HPC/2(1)/2014/ CORP:DG:3191) grant to DM. SP was supported by research fellowship from National Institute of Immunology, New Delhi.

## Conflicts of Interest

The authors declare no conflicts of interest.

## Notes

**Funding statement:** This work was supported by the Department of Biotechnology, Government of India grant to National Institute of Immunology, New Delhi. DM acknowledges financial support from Department of Biotechnology (DBT), India under R&D project grants (BICB: BT/PR40325/BTIS/137/1/2020), (DIAU: BT/BI/TCB/007/2021) and (NNP: BT/PR40267/BTIS/137/67/2023 & BT/PR40160/BTIS/137/64/2023). The work was also supported by National Supercomputing Mission, MeiTY, India under NSM-NPGDD project (MeitY/R&D/HPC/2(1)/2014/ CORP:DG:3191) grant to DM. SP was supported by research fellowship from National Institute of Immunology, New Delhi.

### Competing Interest Statement

The authors have declared no competing interest.

## REFERENCES

1. Yarden, Y. and M.X. Sliwkowski, Untangling the ErbB signalling network. Nat Rev Mol Cell Biol, 2001. 2(2): p. 127–37.

2. Levantini, E., et al., EGFR signaling pathway as therapeutic target in human cancers. Semin Cancer Biol, 2022. 85: p. 253–275.

3. Hanif, F., et al., Glioblastoma Multiforme: A Review of its Epidemiology and Pathogenesis through Clinical Presentation and Treatment. Asian Pac J Cancer Prev, 2017. 18(1): p. 3–9.

4. Sok, J.C., et al., Mutant epidermal growth factor receptor (EGFRvIII) contributes to head and neck cancer growth and resistance to EGFR targeting. Clin Cancer Res, 2006. 12(17): p. 5064–73.

5. Harrison, P.T., S. Vyse, and P.H. Huang, Rare epidermal growth factor receptor (EGFR) mutations in non-small cell lung cancer. Semin Cancer Biol, 2020. 61: p. 167–179.

6. Xu, M.J., D.E. Johnson, and J.R. Grandis, EGFR-targeted therapies in the post-genomic era. Cancer Metastasis Rev, 2017. 36(3): p. 463–473.

7. Xiong, H.Q., et al., Cetuximab, a monoclonal antibody targeting the epidermal growth factor receptor, in combination with gemcitabine for advanced pancreatic cancer: a multicenter phase II Trial. J Clin Oncol, 2004. 22(13): p. 2610–6.

8. Das, D., L. Xie, and J. Hong, Next-generation EGFR tyrosine kinase inhibitors to overcome C797S mutation in non-small cell lung cancer (2019-2024). RSC Med Chem, 2024. 15(10): p. 3371–94.

9. Schultz, D.F., D.D. Billadeau, and S.D. Jois, EGFR trafficking: effect of dimerization, dynamics, and mutation. Front Oncol, 2023. 13: p. 1258371.

10. Tan, J., et al., Electron transfer-triggered imaging of EGFR signaling activity. Nat Commun, 2022. 13(1): p. 594.

11. Laudadio, E., L. Mangano, and C. Minnelli, Chemical Scaffolds for the Clinical Development of Mutant-Selective and Reversible Fourth-Generation EGFR-TKIs in NSCLC. ACS Chem Biol, 2024. 19(4): p. 839–854.

12. Shan, Y., et al., Transitions to catalytically inactive conformations in EGFR kinase. Proc Natl Acad Sci U S A, 2013. 110(18): p. 7270–5.

13. Modi, V. and R.L. Dunbrack, Jr., Defining a new nomenclature for the structures of active and inactive kinases. Proc Natl Acad Sci U S A, 2019. 116(14): p. 6818–6827.

14. Zhang, X., et al., An allosteric mechanism for activation of the kinase domain of epidermal growth factor receptor. Cell, 2006. 125(6): p. 1137–49.

15. Park, J.H., et al., Erlotinib binds both inactive and active conformations of the EGFR tyrosine kinase domain. Biochem J, 2012. 448(3): p. 417–23.

16. Jia, Y., et al., Overcoming EGFR(T790M) and EGFR(C797S) resistance with mutant-selective allosteric inhibitors. Nature, 2016. 534(7605): p. 129–32.

17. Planken, S., et al., Discovery of N-((3R,4R)-4-Fluoro-1-(6-((3-methoxy-1-methyl-1H-pyrazol-4-yl)amino)-9-methyl-9H-purin-2-yl)pyrrolidine-3-yl)acrylamide (PF-06747775) through Structure-Based Drug Design: A High Affinity Irreversible Inhibitor Targeting Oncogenic EGFR Mutants with Selectivity over Wild-Type EGFR. J Med Chem, 2017. 60(7): p. 3002–3019.

18. Galdadas, I., et al., Structural basis of the effect of activating mutations on the EGF receptor. Elife, 2021. 10.

19. Songtawee, N., D.R. Bevan, and K. Choowongkomon, Molecular dynamics of the asymmetric dimers of EGFR: simulations on the active and inactive conformations of the kinase domain. J Mol Graph Model, 2015. 58: p. 16–29.

20. Zhao, W., et al., Rare mutation-dominant compound EGFR-positive NSCLC is associated with enriched kinase domain-resided variants of uncertain significance and poor clinical outcomes. BMC Med, 2023. 21(1): p. 73.

21. Borgeaud, M., et al., Unveiling the Landscape of Uncommon EGFR Mutations in NSCLC-A Systematic Review. J Thorac Oncol, 2024. 19(7): p. 973–983.

22. Yun, C.H., et al., The T790M mutation in EGFR kinase causes drug resistance by increasing the affinity for ATP. Proc Natl Acad Sci U S A, 2008. 105(6): p. 2070–5.

23. Wang, S., S. Cang, and D. Liu, Third-generation inhibitors targeting EGFR T790M mutation in advanced non-small cell lung cancer. J Hematol Oncol, 2016. 9: p. 34.

24. To, C., et al., Single and Dual Targeting of Mutant EGFR with an Allosteric Inhibitor. Cancer Discov, 2019. 9(7): p. 926–943.

25. Jang, J., et al., Mutant-Selective Allosteric EGFR Degraders are Effective Against a Broad Range of Drug-Resistant Mutations. Angew Chem Int Ed Engl, 2020. 59(34): p. 14481–14489.

26. Pal, S., A. Pal, and D. Mohanty, SG-ML-PLAP: A structure-guided machine learning-based scoring function for protein-ligand binding affinity prediction. Protein Sci, 2025. 34(1): p. e5257.

27. Pettersen, E.F., et al., UCSF Chimera--a visualization system for exploratory research and analysis. J Comput Chem, 2004. 25(13): p. 1605–12.

28. Webb, B. and A. Sali, Comparative Protein Structure Modeling Using MODELLER. Curr Protoc Bioinformatics, 2016. 54: p. 5 6 1–5 6 37.

29. Jakalian, A., D.B. Jack, and C.I. Bayly, Fast, efficient generation of high-quality atomic charges. AM1-BCC model: II. Parameterization and validation. J Comput Chem, 2002. 23(16): p. 1623–41.

30. Wang, J., et al., Automatic atom type and bond type perception in molecular mechanical calculations. J Mol Graph Model, 2006. 25(2): p. 247–60.

31. Wang, J., et al., Development and testing of a general amber force field. J Comput Chem, 2004. 25(9): p. 1157–74.

32. D.A. Case, K.B., I.Y. Ben-Shalom, S.R. Brozell, D.S. Cerutti, T.E. Cheatham, III, V.W.D. Cruzeiro,, et al., AMBER 2020, University of California, San Francisco. 2020.

33. Maier, J.A., et al., ff14SB: Improving the Accuracy of Protein Side Chain and Backbone Parameters from ff99SB. J Chem Theory Comput, 2015. 11(8): p. 3696–713.

34. Roe, D.R. and T.E. Cheatham, 3rd, PTRAJ and CPPTRAJ: Software for Processing and Analysis of Molecular Dynamics Trajectory Data. J Chem Theory Comput, 2013. 9(7): p. 3084–95.

35. Hunter, J.D., Matplotlib: A 2D graphics environment. Computing in Science & Engineering, 2007. 9(3): p. 90–95.

36. Genheden, S. and U. Ryde, The MM/PBSA and MM/GBSA methods to estimate ligand-binding affinities. Expert Opin Drug Discov, 2015. 10(5): p. 449–61.

37. Morris, G.M., et al., AutoDock4 and AutoDockTools4: Automated docking with selective receptor flexibility. J Comput Chem, 2009. 30(16): p. 2785–91.

38. O’Boyle, N.M., et al., Open Babel: An open chemical toolbox. J Cheminform, 2011. 3: p. 33.

39. Ravindranath, P.A., et al., AutoDockFR: Advances in Protein-Ligand Docking with Explicitly Specified Binding Site Flexibility. PLoS Comput Biol, 2015. 11(12): p. e1004586.

40. Eberhardt, J., et al., AutoDock Vina 1.2.0: New Docking Methods, Expanded Force Field, and Python Bindings. J Chem Inf Model, 2021. 61(8): p. 3891–3898.

41. Kannan, S., et al., Conformational landscape of the epidermal growth factor receptor kinase reveals a mutant specific allosteric pocket. Chem Sci, 2018. 9(23): p. 5212–5222.

42. Sutto, L. and F.L. Gervasio, Effects of oncogenic mutations on the conformational free-energy landscape of EGFR kinase. Proc Natl Acad Sci U S A, 2013. 110(26): p. 10616–21.

43. Xiong, G., et al., ADMETlab 2.0: an integrated online platform for accurate and comprehensive predictions of ADMET properties. Nucleic Acids Res, 2021. 49(W1): p. W5–W14.

