## Supplementary Information for "Design of Allosteric Inhibitors for Mutant EGFR by Combined use of Machine Learning and Molecular Dynamics Simulations"

**Bioinformatics Center**

**BRIC-National Institute of Immunology**

**Aruna Asaf Ali Marg, New Delhi – 110067, India**

^1^Bioinformatics Center, BRIC-National Institute of Immunology, New Delhi, India

**^*^Address correspondence to:**

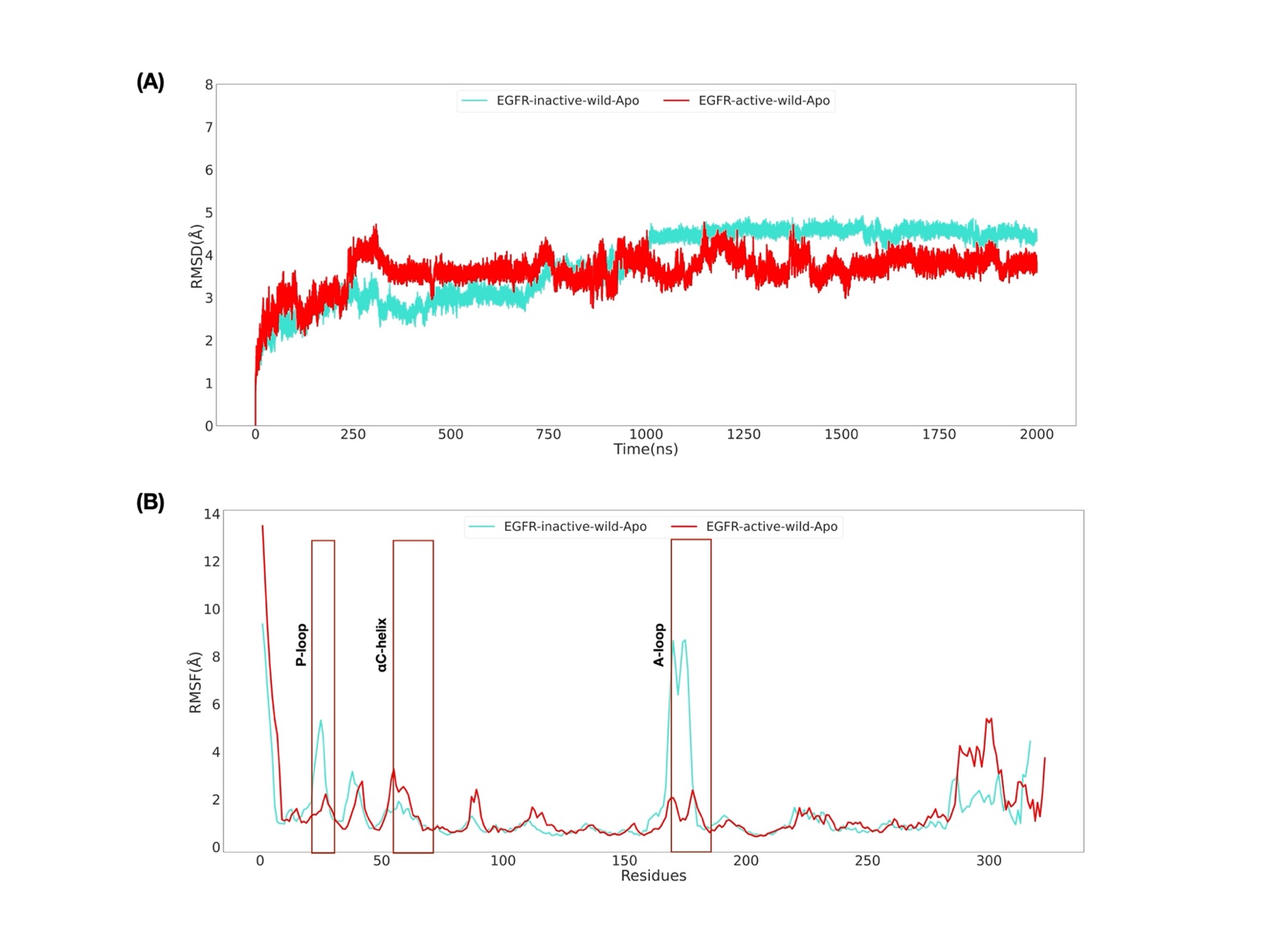

**Supplementary Figure S1.** RMSD (A) and RMSF (B) plot of active and inactive wild-type apo-EGFR upon 2µs simulation. ***Note:** Residue numbering in this figure reflects the modelled EGFR kinase domain (residues 1-317), which corresponds to residues beginning at position 698 of the full-length EGFR protein.

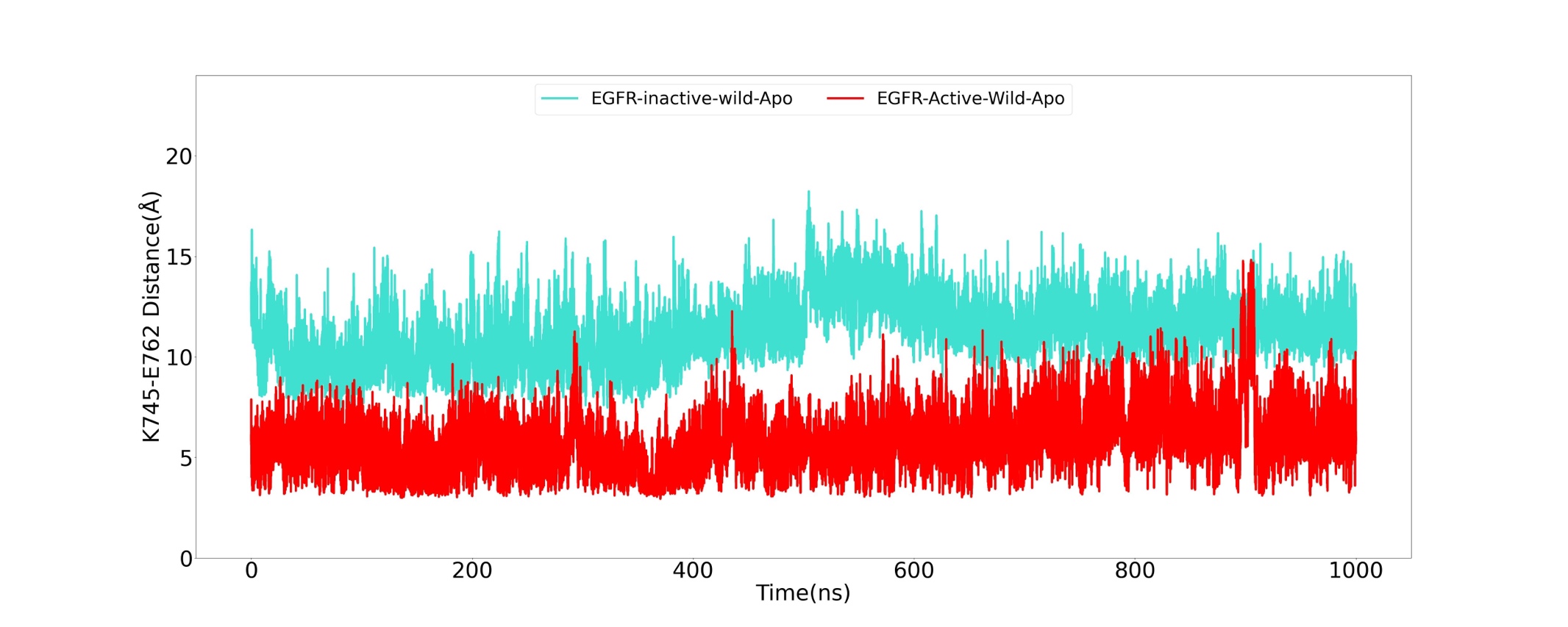

**Supplementary Figure S2.** Time series plot of the distance between NZ and CD atom of K745 and E762 salt bridge forming residues, respectively, for the simulation done using 13Å a non-bonded cutoff distance. Distance calculated for the conformations observed during 1 µs simulations of active and inactive wild-type apo-EGFR.

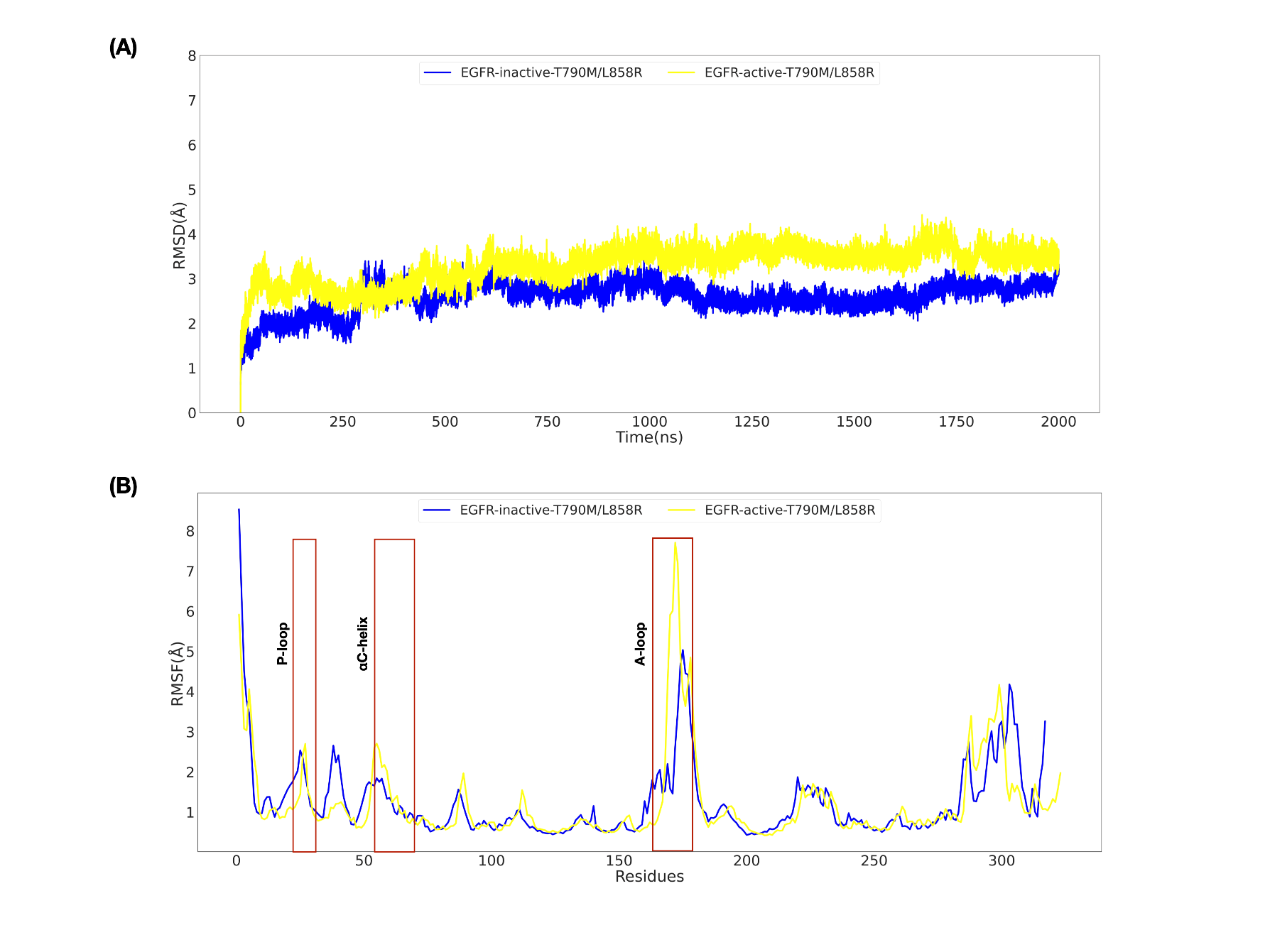

**Supplementary Figure S3.** RMSD (A) and RMSF (B) plot of active and inactive apo-EGFR^L858R/T790M^ upon 2µs simulation. ***Note:** Residue numbering in this figure reflects the modelled EGFR kinase domain (residues 1-317), which corresponds to residues beginning at position 698 of the full-length EGFR protein.

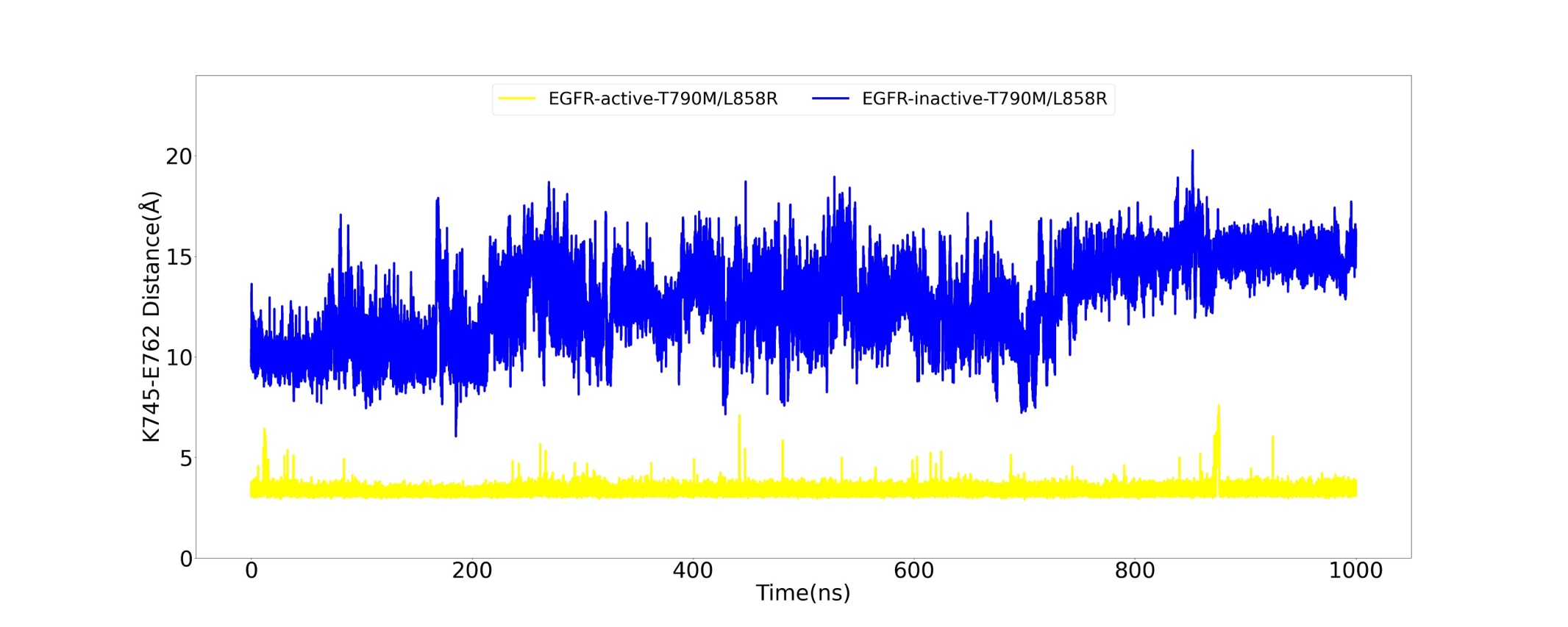

**Supplementary Figure S4.** Time series plot of the distance between NZ and CD atom of K745 and E762 salt bridge forming residues, respectively, for the simulation done using 13Å, a non-bonded cutoff distance. Distance calculated for the conformations observed during 1 µs simulations of active and inactive apo-EGFR^L858R/T790M^.

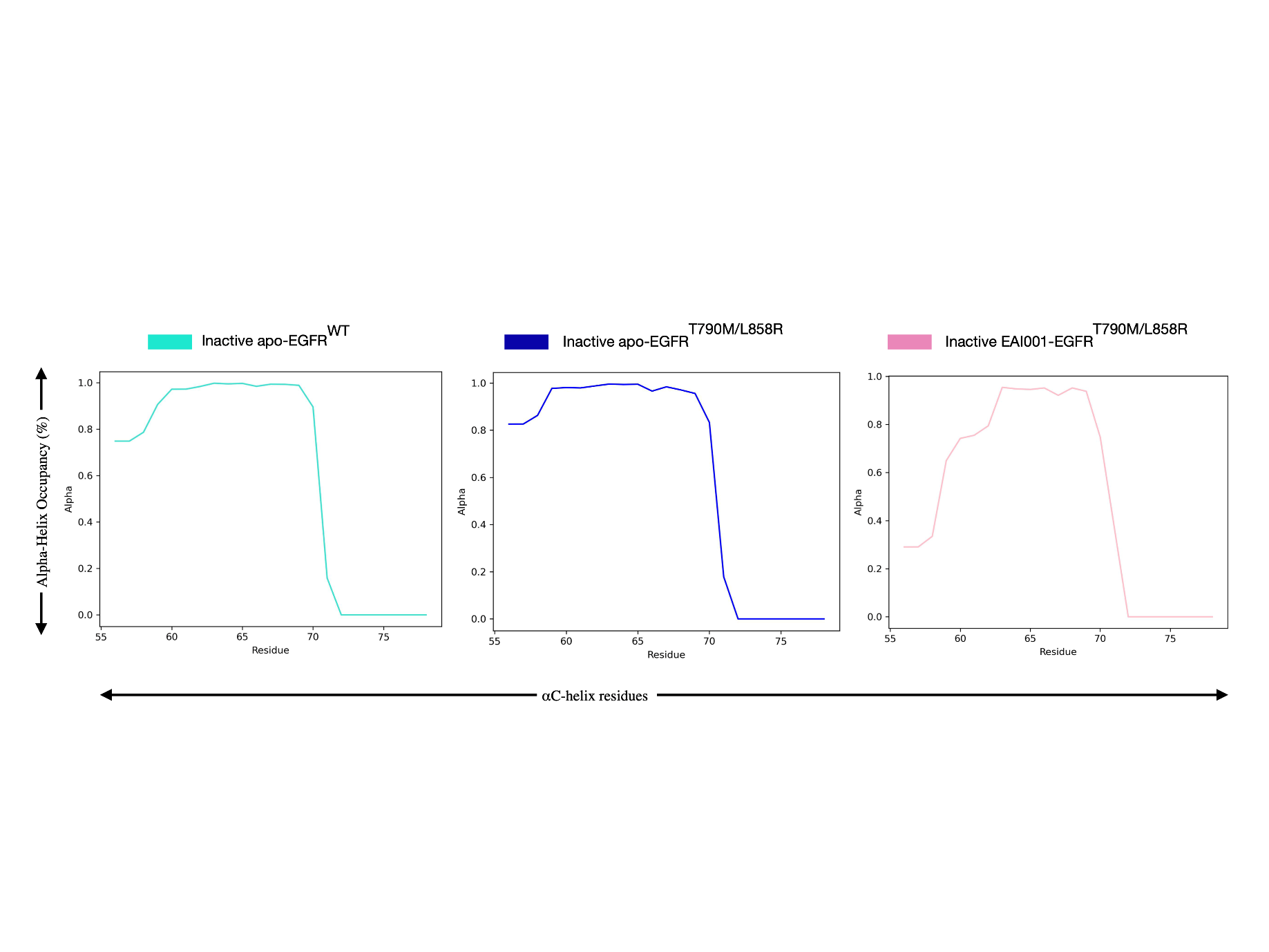

**Supplementary Figure S5.** Secondary structure analysis: Alpha-helix occupancy of αC-helix residues. The plot displays the percentage of simulation time that each αC-helix residue spends in an alpha-helical conformation across 2 µs simulations for inactive apo-EGFR^Wild^, apo−EGFR^L858R/T790M^, and EAI001−EGFR^L858R/T790M^.

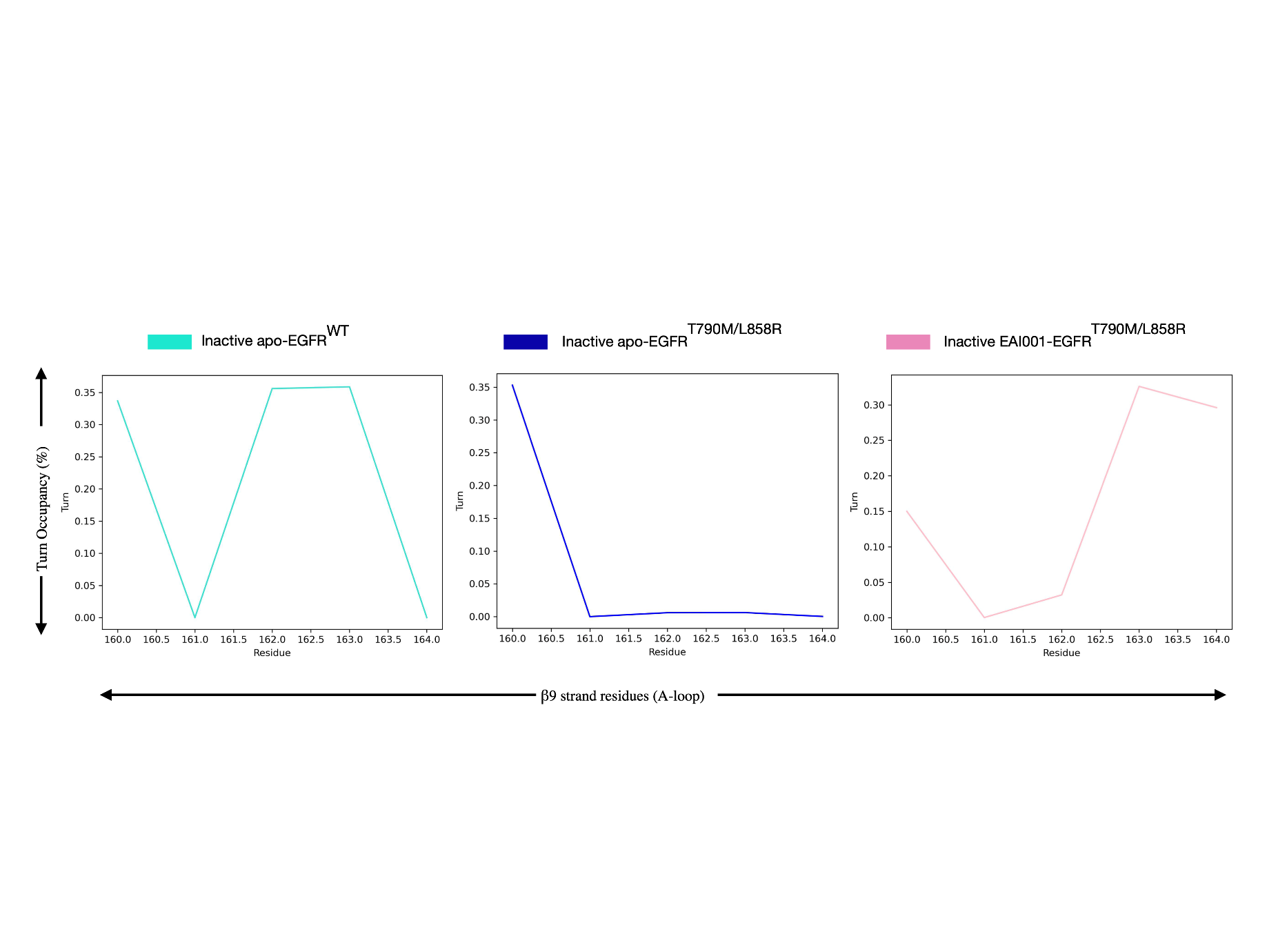

**Supplementary Figure S6.** Secondary structure analysis: Turns occupancy of β9 strand residues of the A-loop. The plot displays the percentage of simulation time each β9 strand residue of the A-loop spends in a turn conformation across 2 µs simulations for inactive apo-EGFR^Wild^, apo−EGFR^L858R/T790M^, and EAI001−EGFR^L858R/T790M^.

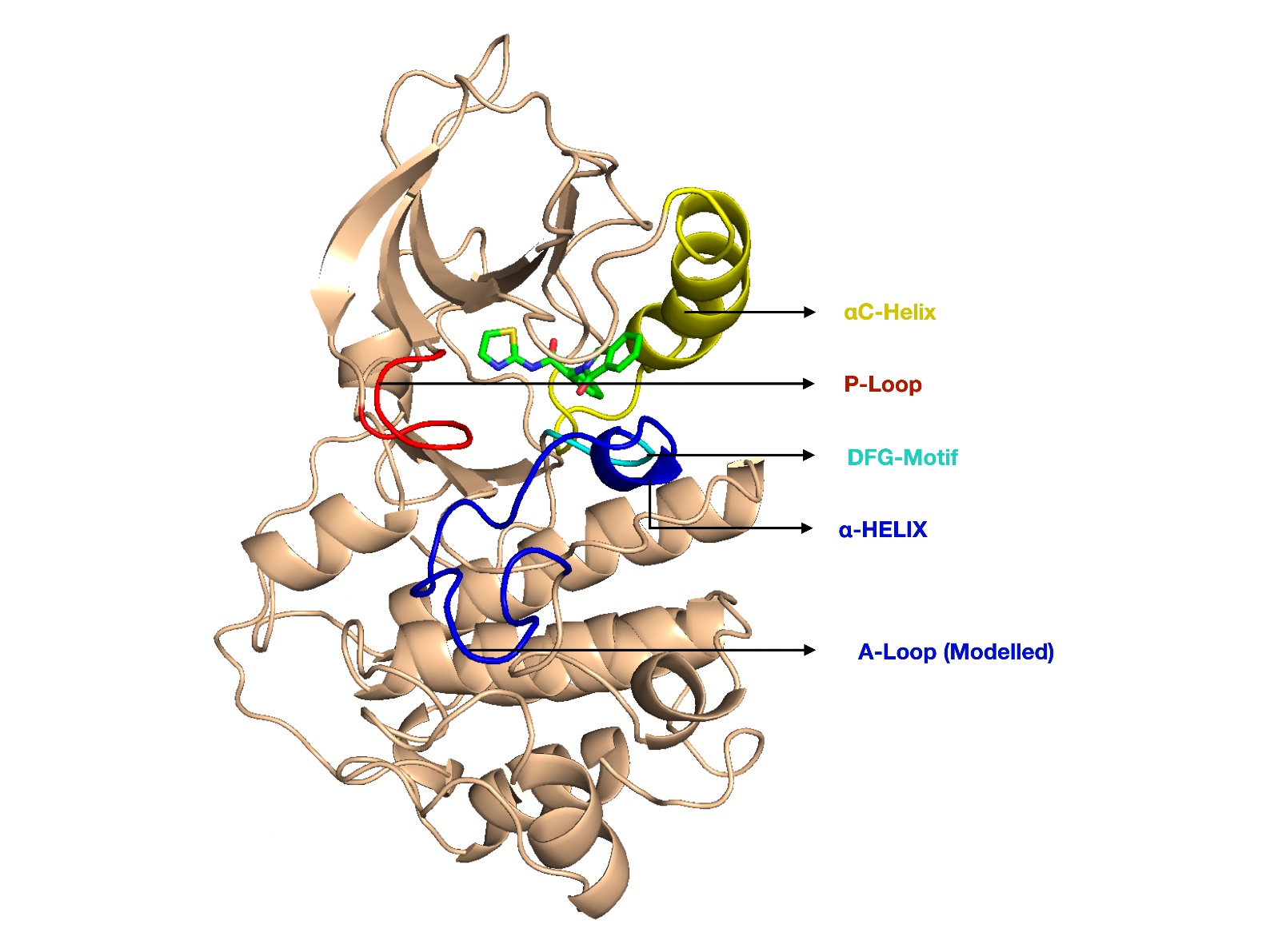

**Supplementary Figure S7.** Conversion of the β9 strand residues into an α-helix at the N-terminal of the A-loop after 8.5 µs.

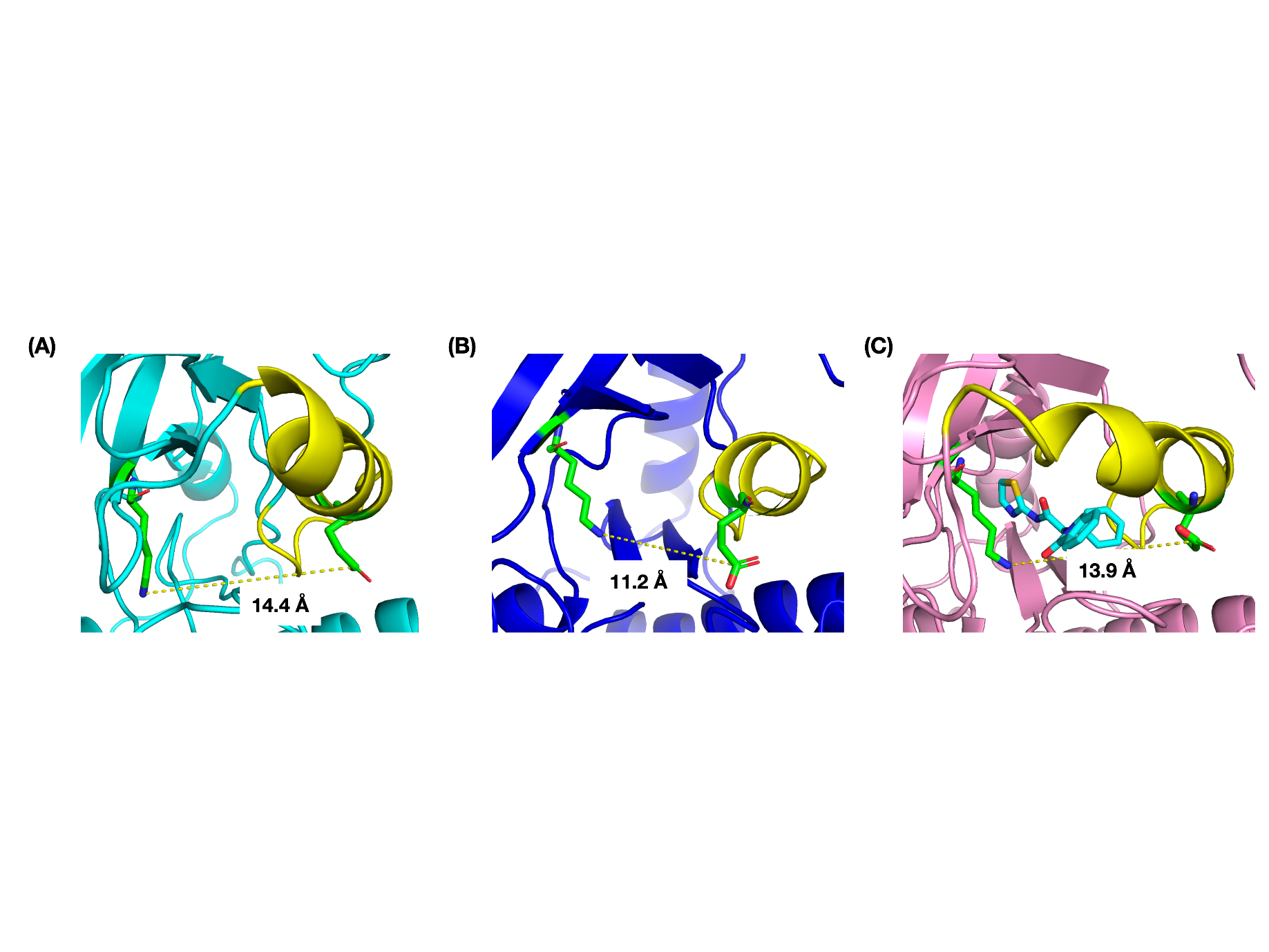

**Supplementary Figure S8.** Salt bridge distances between K745(NZ-atom) and E762(CD-atom) observed in representative conformations extracted from highly populated clusters in the PCA plot (Panels A, B, and C of Figure 7) for inactive apo-EGFR^Wild^, apo−EGFR^L858R/T790M^, and EAI001−EGFR^L858R/T790M^.

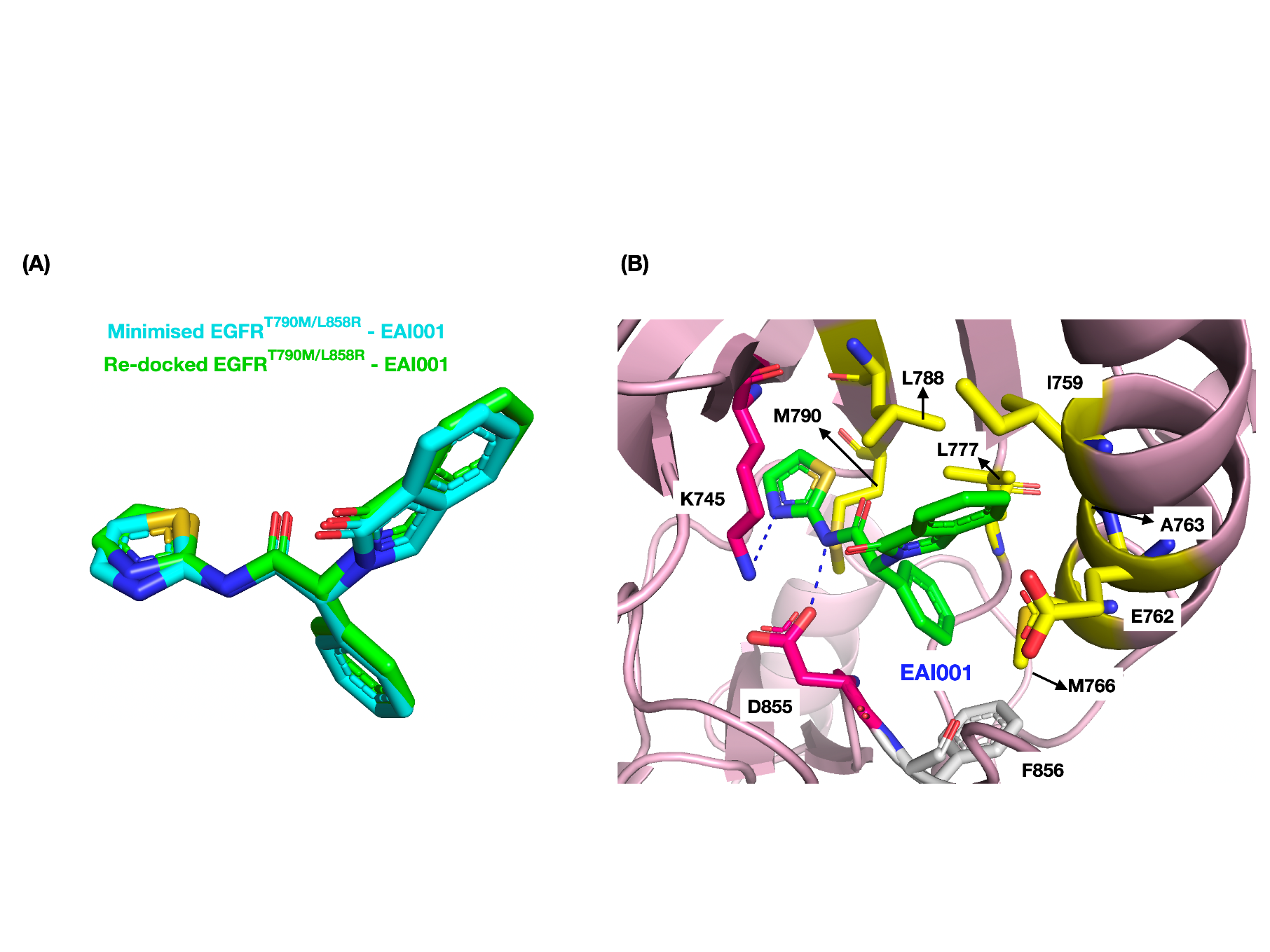

**Supplementary Figure S9.** Validation of docking parameters via re-docking of minimised EGFR^L858R/T790M^ in complex with EAI001 inhibitor. Superimposed image of minimised and re-docked structure of EAI001 on the allosteric pocket of EGFR^L858R/T790M^ (A). Hydrophobic (yellow) and hydrogen bond (pink) interacting residue of EGFR^L858R/T790M^ allosteric pocket with the re-docked ligand (B).

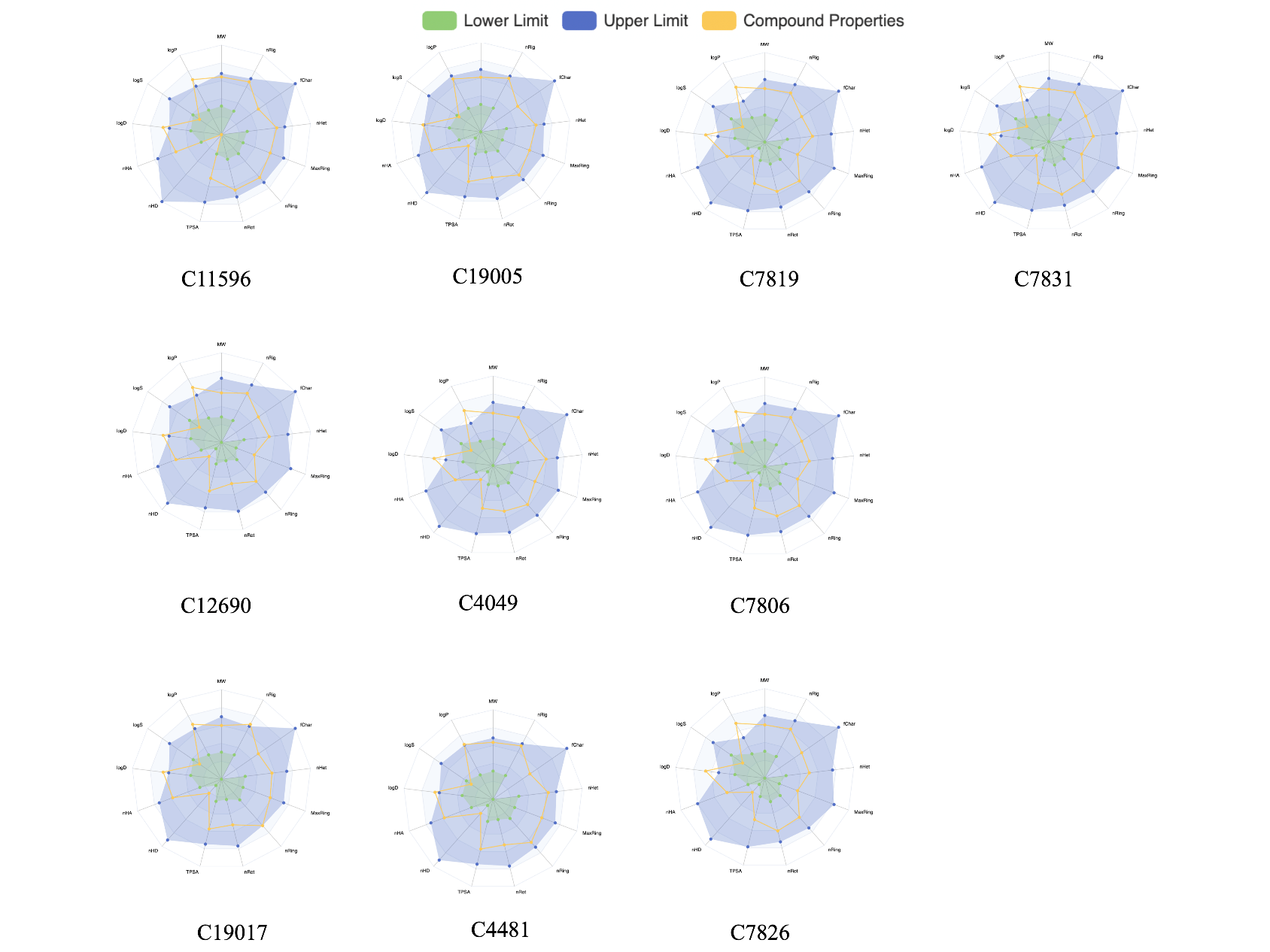

**Supplementary Figure S10.** Radar chart of drug likeliness and physicochemical properties of the top ten selected drugs predicted using the ADMETlab 3.0 software.

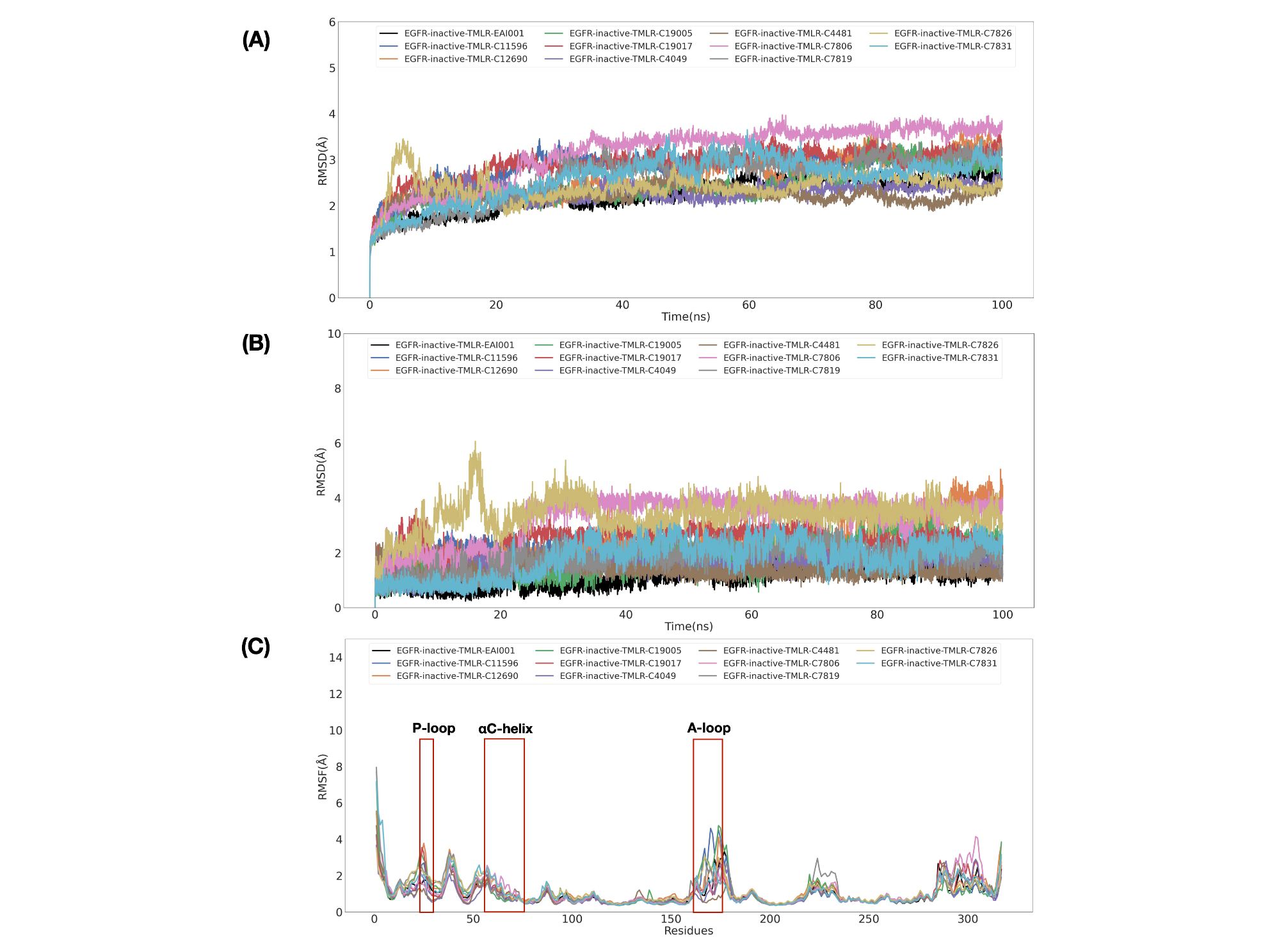

**Supplementary Figure S11.** Trajectory analysis of simulations of top ten compounds with EGFR^L858R/T790M^. RMSD plot of EGFR^L858R/T790M^ when simulated with EAI001 and selected compounds (A). RMSD plot EAI001 and selected compounds after the simulation (B). RMSF plot of EGFR^L858R/T790M^ residues after the simulation (C). ***Note:** Residue numbering in (**C**) reflects the modelled EGFR kinase domain (residues 1-317), which corresponds to residues beginning at position 698 of the full-length EGFR protein.

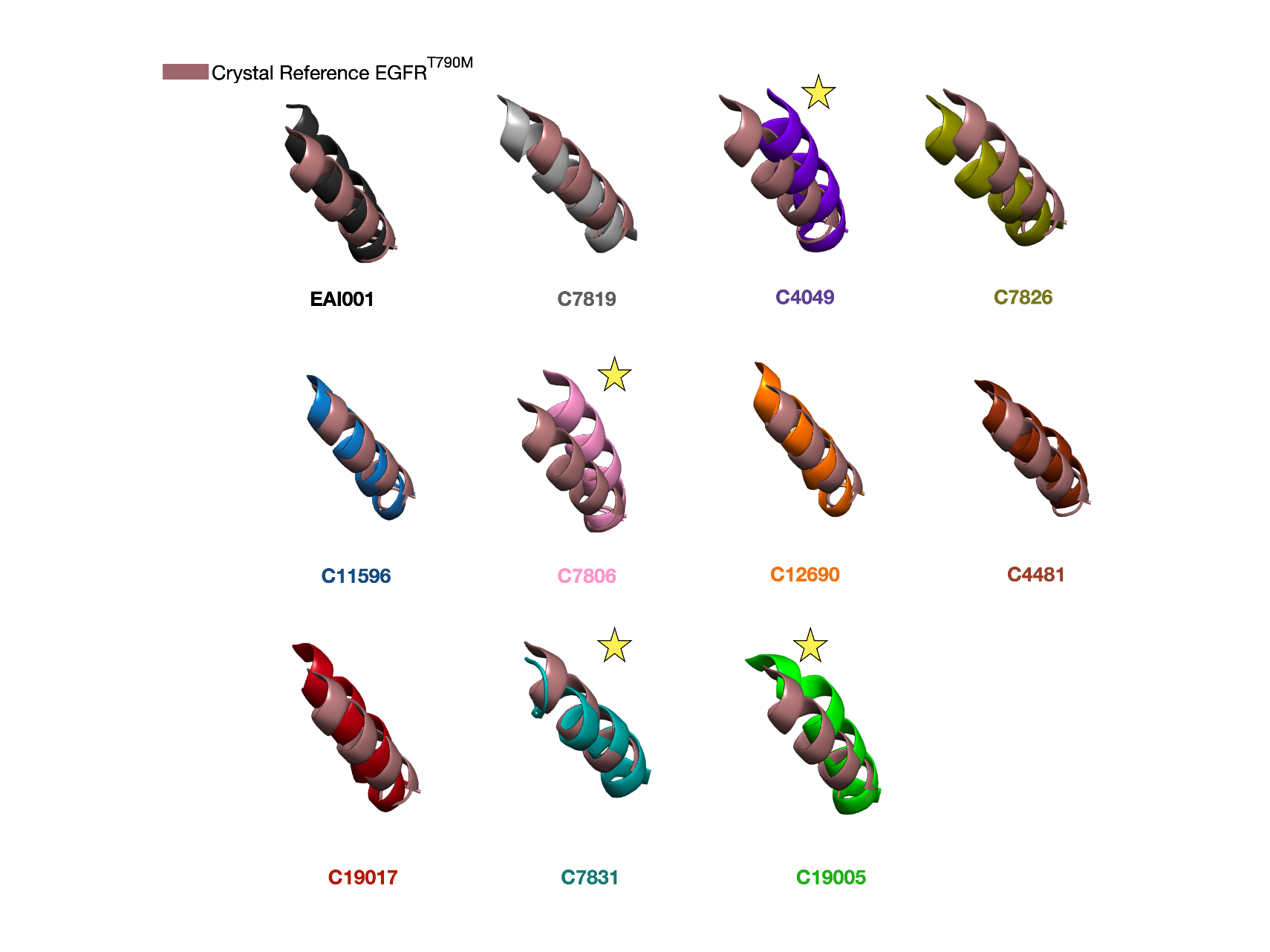

**Supplementary Figure S12.** Superimposition of the EGFRL858R/T790M kinase conformations showing αC-helix (taken at the end of 100ns simulation with top ten compounds) onto the inactive EGFR crystal structure (PDB ID: 5D41) taken as a reference.
